# The UBAP2L ortholog PQN-59 contributes to stress granule assembly and development in *C. elegans*

**DOI:** 10.1101/2021.04.16.440123

**Authors:** Simona Abbatemarco, Alexandra Bondaz, Francoise Schwager, Jing Wang, Christopher M Hammell, Monica Gotta

## Abstract

When exposed to stressful conditions, eukaryotic cells respond by inducing the formation of cytoplasmic ribonucleoprotein complexes called stress granules. Stress granules are thought to have a protective function but their exact role is still unclear. Here we use *C. elegans* to study two proteins that have been shown to be important for stress granule assembly in human cells: PQN-59, the ortholog of human UBAP2L, and GTBP-1, the ortholog of the human G3BP1 and G3BP2 proteins. Both proteins fall into stress granules in the embryo and in the germline when *C. elegans* is exposed to stressful conditions. None of the two proteins is essential for the assembly of stress induced granules, but the granules formed in absence of PQN-59 or GTBP-1 are less numerous and dissolve faster than the ones formed in control embryos. Despite these differences, *pqn-59* or *gtbp-1* mutant embryos do not show a higher sensitivity to stress than control embryos. *pqn-59* mutants display reduced progeny and a high percentage of embryonic lethality, phenotypes that are not dependent on stress exposure and that are not shared with *gtbp-1* mutants. Our data indicate that both GTBP-1 and PQN-59 contribute to stress granule formation but that PQN-59 is, in addition, required for *C. elegans* development.

**Author summary:** The formation of so-called stress granules is an adaptive response that cells and organisms put into action to cope with changes in internal and environmental conditions and thus to survive to stressful conditions. Although it is generally thought that stress granule formation protects cells from stress-related damage, the exact role of stress granules in cells and organisms is not well understood. Moreover, the mechanisms governing stress granule assembly, and if and how the ability to form stress granules is important for *C. elegans* development is still unclear.

Our work focuses on two conserved proteins, known to be involved in stress granule assembly in mammalian cells, and investigates their role in *C. elegans* embryos. We find that these proteins are important but not essential to assemble stress-induced granules in *C. elegans*. We moreover did not observe a different sensitivity to stress exposure between wild-type and mutant developing embryos, suggesting that at least in these conditions these proteins do not exert a protective role.

## Introduction

Eukaryotic cells are sensitive to changes in internal or environmental parameters, including variations in oxygen supply, salt concentration, pH, temperature or viral infection. Each one of these conditions might be sensed as a stressful stimulus by the cell. In return, cells activate the integrated stress response pathway which leads to translation inhibition of most mRNAs and to the assembly of stress granules (1). Stress granules are membraneless organelles formed by the condensation of proteins and RNA molecules into liquid droplets through a mechanism of liquid-liquid phase separation (2). Different protein entities and RNA molecules are recruited into stress granules and their composition varies according to the cell type and the triggering stress (3,4).

Formation of stress-induced granules is a reversible process, hence removal of the stress stimulus results in dissolution of the granules. The current model describing the pathway through which cells assemble stress granules involves disassembly of the polysomes with consequent translation inhibition either via phosphorylation of the translation initiation factor eIF2α (Eukaryotic Initiation Factor 2 alpha) (5) or via the inhibition of eIF4G (Eukaryotic Initiation Factor 2 G) (6). The mRNAs released from the polysomes are then bound to RNA binding proteins and recruited into the stress granules (7,8). In mammalian cells, together with the translation initiation factor eIF2α, other proteins are important nucleators of stress granules. These include G3BP1 and G3BP2 (Ras GTPase-activating protein-binding protein 1 and 2) and UBAP2L (Ubiquitin Associated Protein 2 Like), which are crucial to drive stress granule assembly in many stress conditions (9–12), and the protein TIA-1 (T-cell-restricted intracellular antigen protein) (5,13).

Although the exact function of stress granules and their importance for cell survival and organismal development have not yet been established, stress granules may exert a protective role on cells when they are exposed to stress (14).

Stress granule assembly and function has been mainly studied in unicellular organisms and cells in culture. The nematode *C. elegans* provides an excellent model to study stress granules and to address their role in organismal viability. The proteins involved in stress granule formation in mammalian cells are conserved and the formation of granules molecularly similar to the mammalian stress granules has been observed in the somatic and germ cells (15–18).

*C. elegans* contains one ortholog of the mammalian G3BP1 and 2, called GTBP-1 (19) and two TIA-1/TIAR orthologs (20), named TIAR-1 and TIAR-2. GTBP-1 has been only recently shown to contribute to stress granule formation in *C. elegans* adult worms (18). TIAR-1 protects germ cells from heat-shock (17) and TIAR-2 granules inhibit axon regeneration (21). The *C. elegans* potential ortholog of UBAP2L is a protein called PQN-59 (Prion-like (glutamine/asparagine-rich) domain bearing protein) (22,23). The similarity between PQN-59 and UBAP2L at the sequence level is only 30% (source: BlastP) but PQN-59 and UBAP2L share a very similar domain organization (Fig 1A). As GTBP-1, PQN-59 is an abundant protein of the entire *C. elegans* proteome (https://pax-db.org/protein/1033201) (24), but its role in *C. elegans* has not been characterized.

**Fig 1.**
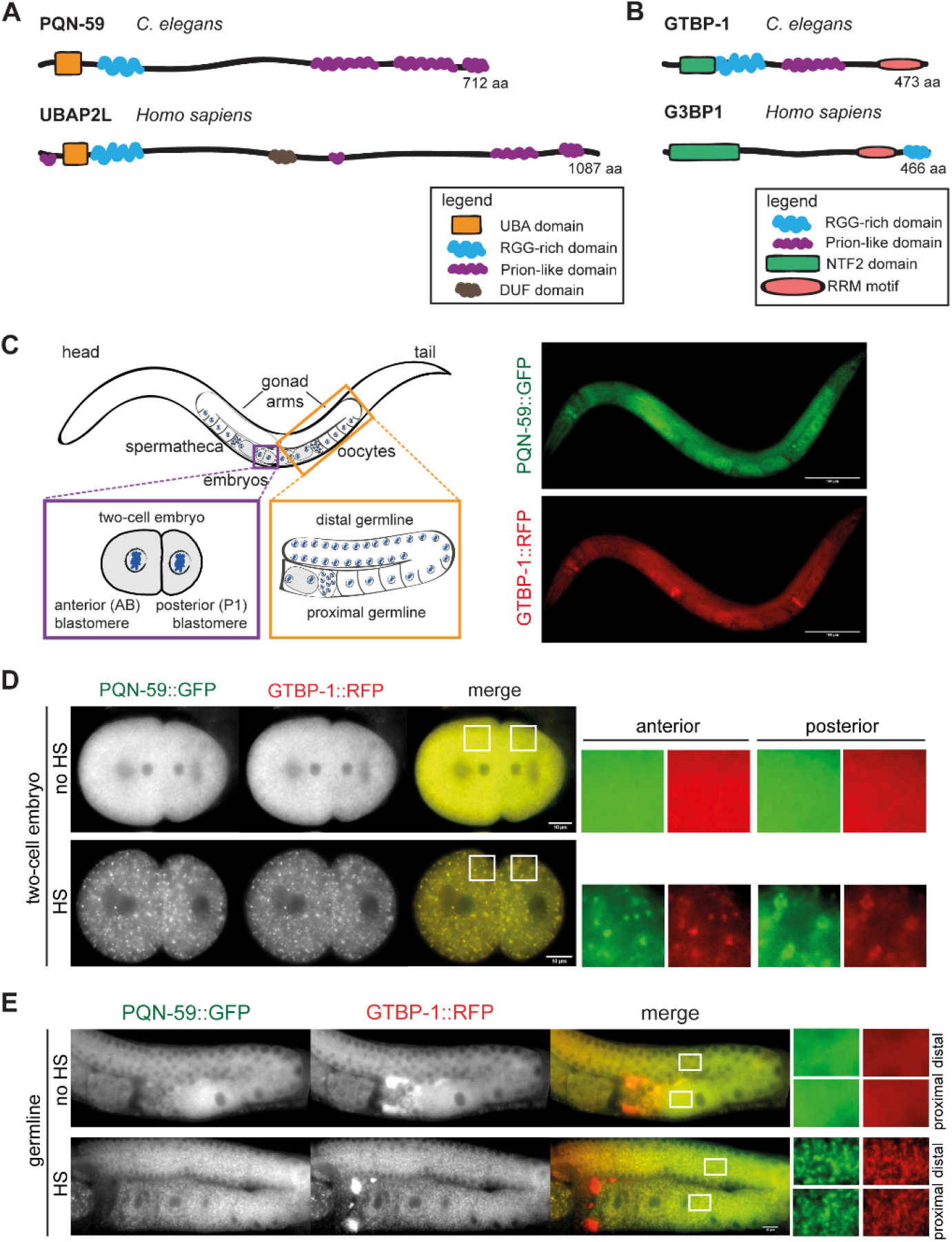
PQN-59 and GTBP-1 co-localize into heat stress-induced granules in the embryo and in the germline. (**A**) and (**B**) Schematic representation of the protein domains of PQN-59 and its human ortholog UBAP2L (**A**) and of GTBP-1 and its human ortholog G3BP1 (**B**). (**C**) Schematic drawing of an adult *C. elegans* worm, with close ups of a two-cell embryo (bottom left, purple square) and of the germline (bottom right, orange square) and images of an adult animal expressing endogenous *pqn-59::GFP;gtbp-1::RFP.* Scale bars represent 100 μm. (**D**) Still frames from time-lapse imaging of *pqn-59::GFP;gtbp-1::RFP* embryos using a CherryTemp temperature-controlled stage. Embryos were imaged at 20°C (no heat-shock, no HS) or at 30°C for 5 min (heat-shock, HS). PQN-59/GTBP-1 granules were observed in 100% of the observed embryos (n=12, N=5). In all figures, white boxes indicate the ROI shown enlarged on the right. For embryos, ROIs are in the anterior AB cell (left) and in the posterior P1 cell (right). (**E**) Germlines of *pqn-59::GFP;gtbp-1::RFP* worms in control conditions (no HS, 20°C) and after heat-stress exposure (HS, 10 min at 35°C). PQN-59/GTBP-1 granules were detected in 98% of the observed gonads (n=51, N=8). In all images of germlines, white boxes in the distal (top) and proximal (bottom) germline show the ROI enlarged on the right. Scale bars represent 10 μm. In all images, ROIs are enlarged 8X (embryos) and 11.5X (germline).

Here we show that different stress stimuli trigger the formation of granules containing both PQN-59 and GTBP-1 in *C. elegans* embryos and germlines. We find that neither of the two proteins is essential for stress granule assembly, but both contribute to this process. However, PQN-59 depletion or deletion results in embryonic lethality and reduced progeny in normal growth condition, phenotypes that are not observed following the depletion or deletion of GTBP-1. This suggests that PQN-59 plays additional roles in the development of worms.

## Results

### PQN-59 is a component of stress granules

The UBAP2L protein is important for stress granule assembly in many stress conditions and acts upstream of the stress granule components G3BP1 and 2 in this process (4,10,12,25). We set out to investigate whether the *C. elegans* ortholog of UBAP2L, called PQN-59 (Fig 1A), is also a component of stress granules.

We used a CRISPR/Cas9 generated strain expressing an endogenous C-terminal fusion of PQN-59 with GFP and of GTBP-1, the ortholog of human G3BP1 and 2 (Fig 1B), with RFP (see Strain List table in Materials and Methods). Both PQN-59 and GTBP-1 are expressed throughout development, and are widely expressed in adult *C. elegans* animals, including the germline and the embryo (Fig 1C, (26)).

Observation of untreated *pqn-59::GFP;gtbp-1::RFP* embryos revealed that both proteins are cytoplasmic (in the embryos and in the germline, Figs 1D and 1E). When embryos were exposed to heat-shock (30°C, 5 minutes) using a temperature-controlled stage, PQN-59 fell into granules in both the anterior and posterior blastomere (Fig 1D). These granules colocalized with GTBP-1 granules (Fig 1D). Similar to stress granules (27), lowering the temperature to 20°C following heat-shock exposure resulted in dissolution of the PQN-59/GTBP-1 granules after about 15 minutes of recovery (S1A Fig and S1 Movie). Staining of untreated wild-type embryos with PQN-59 antibodies confirmed the cytoplasmic localization observed with the GFP CRISPR strain (S1B Fig). The immunostaining signal was abolished after PQN-59 depletion by RNA interference (S1B Fig), confirming the specificity of the antibody. Staining of embryos exposed to heat shock revealed the accumulation of PQN-59 into cytoplasmic granules (S1C Fig), similar to what we observed with the *pqn-59::GFP* strain.

We then asked whether following high temperature exposure PQN-59 and GTBP-1 also fall into granules in the *C. elegans* germline. We found that in *pqn-59::GFP;gtbp-1::RFP* worms exposed to 35°C for 10 minutes, PQN-59 fell into granules in both the distal and proximal germline (Fig 1E). These granules colocalized with GTBP-1 granules (Fig 1E) and dissolved after 10 minutes of incubation at 20°C (S2A Fig), confirming that their formation depends on stress exposure and is reversible. Therefore, heat-stress induces the formation of PQN-59/GTBP-1 containing granules also in the *C. elegans* germline.

We then asked whether PQN-59 falls into granules when worms are exposed to other stresses. Sodium Arsenite induces oxidative stress triggering the formation of stress granules (16). Adult worms incubated in a solution containing 20 mM Arsenite for 5 hours displayed granules containing both PQN-59 and GTBP-1 (S2B Fig). The granules were observed both in the proximal and distal germline.

The translation inhibitor Puromycin promotes polysome disassembly and stress granule formation (17). We therefore asked whether incubation with Puromycin would induce PQN-59 and GTBP-1 granule formation. As shown in S2B Fig, worms incubated for 4 hours in a solution containing 10 mg/ml of Puromycin showed the appearance of PQN-59 granules that colocalized with GTBP-1 in the distal and the proximal germline.

To conclude, the exposure of *C. elegans* animals to heat-shock, Arsenite and Puromycin, results in the formation of PQN-59 cytoplasmic granules that colocalize with the known stress granule component GTBP-1. The granules are reversible as they dissolve when the stress is removed. These data indicate that PQN-59 is a stress granule component.

### PQN-59 is important for the formation of stress-induced GTBP-1 granules

We next asked whether PQN-59 is required to form stress granules in the embryo and in the germline.

In *ctrl(RNAi)* embryos exposed to 34°C prior to fixation, both PQN-59 and GTBP-1 fell into granules (Fig 2A). When heat-shock was applied to *pqn-59(RNAi)* embryos, GTBP-1 fell into granule in both the anterior and posterior cells. However, whereas in heat-shocked *ctrl(RNAi)* embryos GTBP-1 granules appeared like small, spherical and defined speckles, in heat-shocked PQN-59-depleted embryos, GTBP-1 formed larger and more diffuse granules (Fig 2A). Quantifications of the GTBP-1 signal revealed that in PQN-59-depleted embryos the number and the intensity of GTBP-1 granules are reduced compared to control embryos (Fig 2B). The depletion of PQN-59 did not result in a change of GTBP-1 levels (S3A and S3B Figs). However, in *pqn-59(RNAi)* embryos that were not exposed to heat-shock, GTBP-1 fell into granules in the posterior P1 blastomere that colocalized with a P body marker (S3A and S3C Figs).

**Fig 2.**
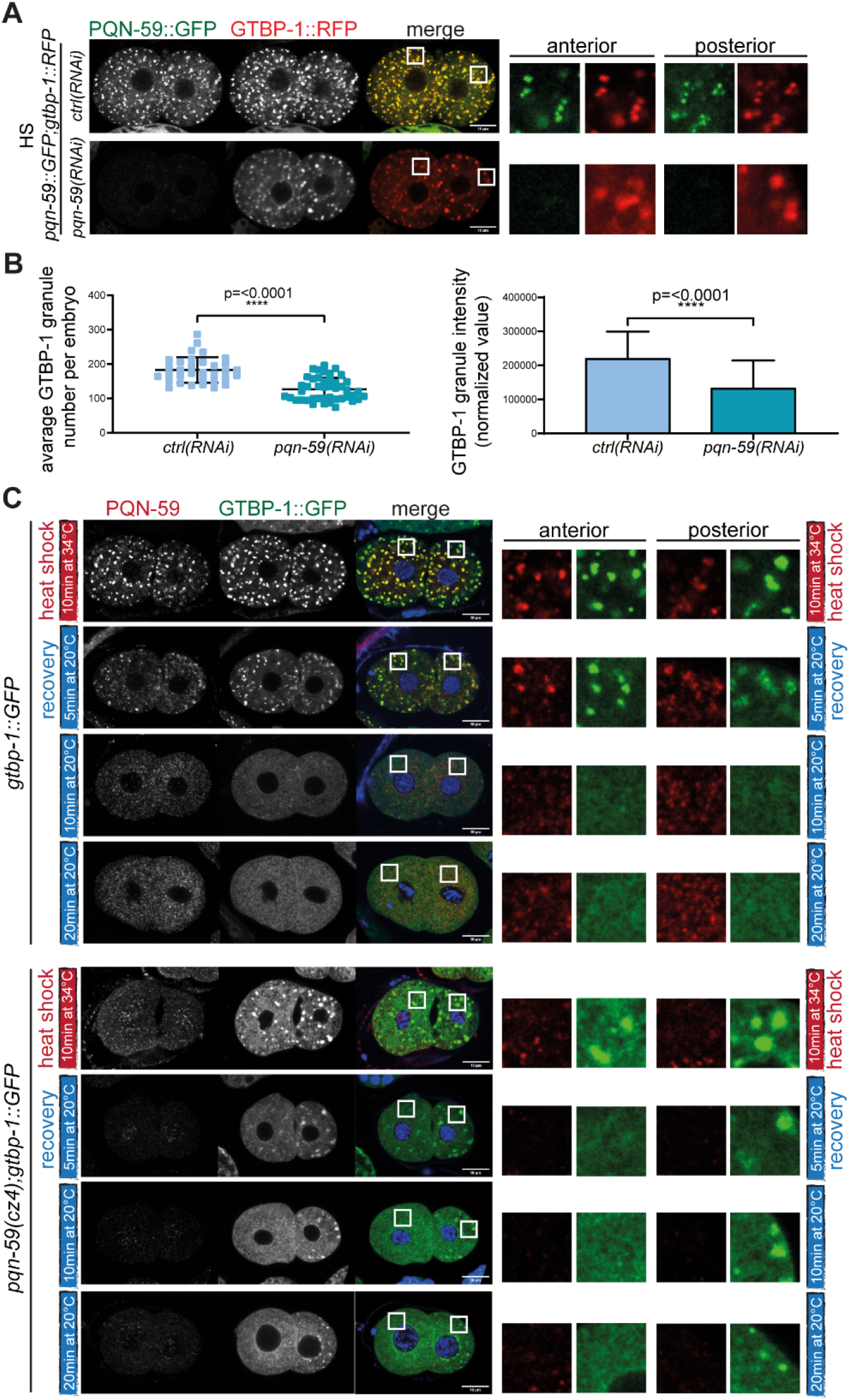
PQN-59 depletion impairs stress-induced GTBP-1 granule formation. (**A**) Single confocal planes of *pqn-59::GFP;gtbp-1::RFP* fixed two-cell embryos treated with the indicated RNAi and exposed to heat-shock (HS, 34°C for 10 minutes) before fixation. (**B**) Quantification of the average GTBP-1 granule number (left) and the average normalized GTBP-1 granule intensity (right) per embryo *(ctrl(RNAi)* n=33; *pqn-59(RNAi)* n=44, N=4). Error bars indicate S.D. The P-value was determined using Student’s t-test. (**C**) Single confocal planes of *gtbp-1::GFP* and *pqn-59(cz4);gtbp-1::GFP* fixed embryos immunostained with PQN-59 antibodies (red). GTBP-1 GFP signal is in green and DNA was counterstained with DAPI (blue). Embryos were fixed at different time points: immediately after heat-shock exposure (10 minutes at 34°C, red vertical line) and after recovery at 20°C for 5, 10, or 20 minutes (blue vertical line). For each time point in **C**, between 15 and 21 embryos were analyzed for the *gtbp-1::GFP* and between 7 and 13 for the *pqn-59(cz4);gtbp-1::GFP* (N=4). Scale bars represent 10 μm. Enlarged ROIs are on the right.

To exclude that a residual pool of PQN-59 after RNAi depletion could account for GTBP-1 granule formation after stress exposure, we inserted a stop codon in the second exon of PQN-59 in a strain expressing GTBP-1::GFP (S3D Fig). As revealed through western blot analysis, expression of PQN-59 was absent in this strain (S3E Fig). Similarly to what we observed with the depletion of PQN-59 in the *pqn-59::GFP;gtbp-1::RFP* strain, GTBP-1 fell into granules in the posterior P1 blastomere in the *pqn-59(cz4)* embryos (S3F Fig). In embryos exposed to heat-shock, GTBP-1 formed large and diffuse aggregates.

As shown above (S1A Fig), heat-induced PQN-59/GTBP-1 granules dissolved when the temperature was shifted back to 20°C. Images of wild-type embryos fixed after heat-shock and after 5, 10 and 20 minutes of recovery at 20°C, confirmed that PQN-59/GTBP-1 stress-induced granules are still present after 5 minutes of recovery, and were not detected after 10 minutes (Fig 2C). In *pqn-59(cz4)* embryos, however, the GTBP-1 stress-induced granules were already dissolved after 5 minutes of recovery in the anterior blastomere, therefore showing a faster dissolution timing compared to the parental strain (Fig 2C). The granules in the posterior blastomere did not dissolve, consistent with the fact that their formation is not dependent on heat-shock exposure (S3F Fig).

The observation that after PQN-59 depletion GTBP-1 localization is affected and that GTBP-1 stress-induced granules are reduced in number and less intense, suggests an interdependence in stress granule formation between these two proteins. We therefore tested whether PQN-59 and GTBP-1 interact. In agreement with data in other model systems (28), PQN-59 interacted with GTBP-1 in Two Hybrid assays, as shown by growth on selective medium of yeast colonies expressing GTBP-1 and PQN-59 (S3G Fig).

We then asked whether GTBP-1 granules can form in the germline when PQN-59 is depleted. As shown in S4A Fig, depletion of PQN-59 abolished the formation of GTBP-1 granules in the oocytes (proximal germline). However, GTBP-1 granules were still observed around the nuclei of the syncytial germline (distal germline), indicating that, similar to the situation in the embryo, GTBP-1 granules can still form, although not throughout the entire germline.

The RGG domain of UBAP2L is crucial to nucleate stress granules in human cells (12,25). We deleted this domain in the *pqn-59::GFP;gtbp-1::RFP* strain (S4B Fig) and tested whether PQN-59ΔRGG could still form granules after heat-shock. As shown in Fig S4C, PQN-59ΔRGG was nucleating granules that colocalized with GTBP-1, similar to the granules formed in the wild-type strain. Quantification of both PQN-59 and GTBP-1 granule number and intensity revealed similar values between the wild-type parental strain and the *pqn-59::ΔRGG::GFP;gtbp-1::RFP* strain (S4D Fig).

Our data show that when PQN-59 is absent, GTBP-1 can still form granules after heat-shock but these granules appear different from the stress granules assembled in the control strain and they dissolve with faster dynamics. The deletion of the RGG domain of PQN-59 alone is not sufficient to impair stress granule assembly, indicating that this domain is not essential in this process in *C. elegans* embryos.

### GTBP-1 contributes to the assembly of stress-induced granules

In mammalian cells, the G3BP proteins are crucial to assemble stress granules in many stress conditions (9,11,29,30). We therefore investigated whether GTBP-1 was required for the assembly of PQN-59 granules in *C. elegans* after heat shock.

We depleted GTBP-1 in *pqn-59::GFP;gtbp-1::RFP* worms, and imaged embryos after heat-shock and fixation. In GTBP-1 depleted embryos at 20°C, PQN-59 was diffused in the cytoplasm, as in *ctrl(RNAi)* embryos (S5A Fig). In *ctrl(RNAi)* heat-shocked embryos we observed numerous granules containing PQN-59 and GTBP-1, in both the anterior AB and posterior P1 cells of two-cell embryos (Fig 3A). After GTBP-1 depletion, some PQN-59 granules were still observed in both AB and P1 cells but were smaller and less defined (Fig 3A). A significant decrease in PQN-59 number and intensity could be quantified in GTBP-1-depleted embryos compared to control ones (Fig 3B). Heat-shock of *gtbp-1(ax2029)* mutant embryos followed by PQN-59 staining resulted in a phenotype similar to the GTBP-1 depletion (S5B Fig).

**Fig 3.**
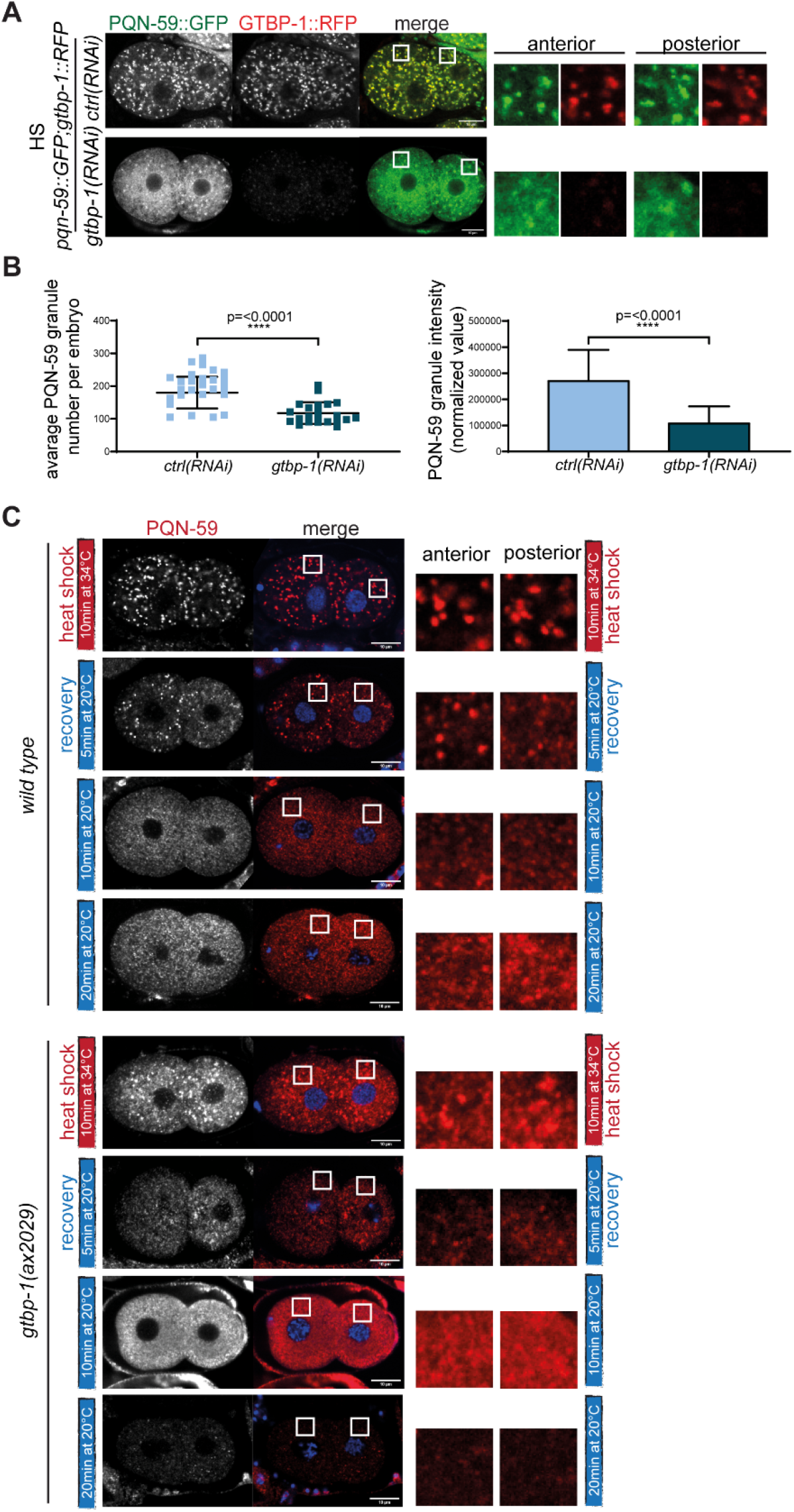
GTBP-1 depletion impairs PQN-59 stress-induced granule formation. (**A**) Single confocal planes of *pqn-59::GFP;gtbp-1::RFP* fixed two-cell embryos treated with the indicated RNAi and exposed to heat-shock (HS, 34°C for 10 minutes) before fixation. (**B**) Quantification of the average PQN-59 granule number (left) and the average normalized PQN-59 granule intensity (right) per embryo *(ctrl(RNAi)* n=40; *gtbp-1(RNAi)* n=22, N=4). Error bars indicate S.D. The P-value was determined using Student’s t-test. (**C**) Single confocal planes of *wild-type* and *gtbp-1(ax2029)* fixed two-cell embryos immunostained with PQN-59 antibodies (red). DNA was counterstained with DAPI (blue). Embryos were fixed at different time points: immediately after heat-shock exposure (10 minutes at 34°C, red vertical line) and after recovery at 20°C for 5, 10, or 20 minutes (blue vertical line). Between 6 and 17 embryos were analyzed for each condition and genotype, N=3. Scale bars represent 10 μm. ROIs are enlarged on the right.

The absence of GTBP-1 did not affect the levels of PQN-59, as detected by immunofluorescence in the PQN-59::GFP tagged strain or after anti-PQN-59 antibody staining (S5A and S5B Figs, and quantifications in S5C and S5D Figs), indicating that the impaired stress granule assembly did not depend on a change in protein amount. The small stress-induced PQN-59 granules formed in absence of GTBP-1 also showed a faster dissolution dynamic following stress ceasing (Fig 3C). While in heat-shocked wild-type embryos granules were still present after 5 minutes of recovery at 20°C and started to dissolve after 10 minutes (Fig 3C), in *gtbp-1(ax2029)* mutant embryos the PQN-59 granules started disappearing already after 5 minutes of recovery at 20°C (Fig 3C). This indicates that the biophysical properties of the granules formed in absence of GTBP-1 are altered compared to control conditions.

Depletion of GTBP-1 also impaired PQN-59 granule assembly in the germline. Dim PQN-59 granules were still visible around the nuclei in the distal germline. In the proximal germline, aberrant PQN-59 aggregates were observed (S5E Fig).

We conclude that when GTBP-1 is depleted, stress-exposed embryos contain less numerous and less intense PQN-59 granules. In GTBP-1 depleted germlines, PQN-59 still falls into granules in the distal and in aberrant aggregates in the proximal germline.

### TIAR-1 granules assemble in embryos depleted of GTBP-1 and PQN-59

Since depleting PQN-59 did not abolish formation of GTBP-1 granules and, vice-versa, depleting GTBP-1 did not abolish the formation of PQN-59 granules after exposure to stress, we asked whether depleting both proteins would result in a defect in the formation of stress granules. To address this question we used as a marker the protein TIAR-1. In *C. elegans*, TIAR-1 accumulates in stress granules in the germline (17,31) and in the intestine of the adult worms (18) following exposure to different stresses.

In the *C. elegans* embryo, TIAR-1 is localized in the cytoplasm and it accumulates in the nuclei and the P granules of the germ precursor cells (S6A Fig and (17,20)). We used a strain expressing TIAR-1::GFP (17) and found that following heat shock of the embryo, TIAR-1 accumulated into stress-induced granules which colocalized with PQN-59 (Fig 4A). Depletion of PQN-59 did not abolish TIAR-1 granule formation after heat-stress exposure (Fig 4A). Our quantifications showed that the number and the intensity of TIAR-1 granules was not significantly different compared to the control (Fig 4B). However, we observed a higher variability in the PQN-59 depleted embryos, consistent with the fact that some embryos appeared to have less granules. In *tiar-1::GFP;gtbp-1(ax2029);ctrl(RNAi)* embryos that were heat-shocked, the majority of TIAR-1 granules were detected in the P1 cell (Fig 4A) but the overall number and intensity of TIAR-1 granules did not appear to be different from the wild-type parental strain (Fig 4B). In this condition, consistently with the result showed in Fig 3B, PQN-59 formed granules in both AB and P1 cells, and these granules colocalized with TIAR-1 granules (Fig 4A). When PQN-59 was depleted in the *tiar-1::GFP;gtbp-1(ax2029)* embryos, TIAR-1 granules were still observed after heat-shock, but their number was reduced compared to control *tiar-1::GFP* embryos (Fig 4B). This suggests that the depletion of both PQN-59 and GTBP-1 proteins is not sufficient to abolish the assembly of TIAR-1 stress-induced granules. In embryos that were not exposed to heat-shock, the localization and appearance of TIAR-1 was not affected by the depletion of PQN-59, the mutation of GTBP-1 or both (S6A Fig).

**Fig 4.**
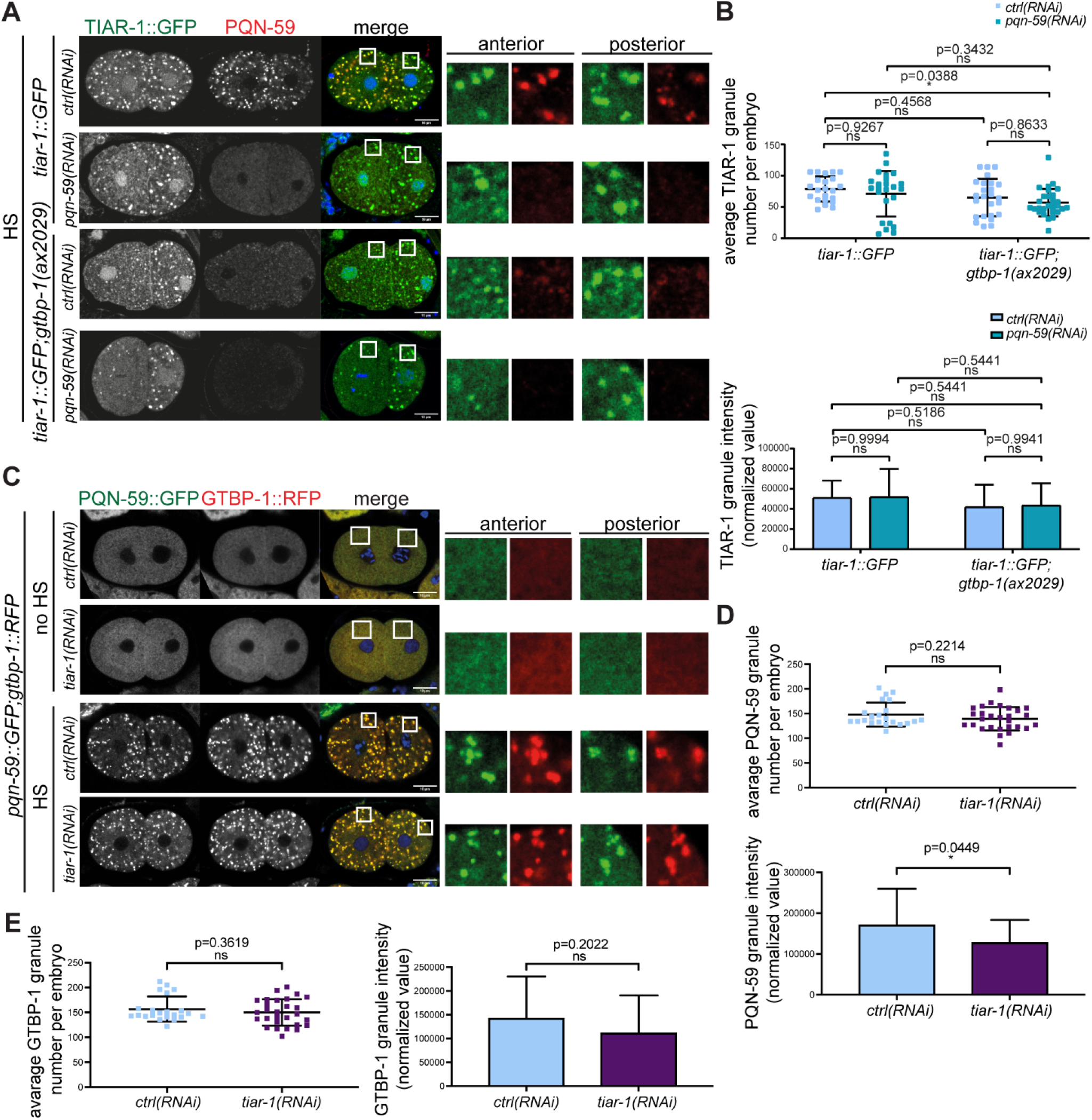
The number of TIAR-1 granules is reduced in *pqn-59(RNAi); gtbp-1(ax2029)* embryos. (**A**) Single confocal planes of *tiar-1::GFP* and *tiar-1::GFP;gtbp-1(ax2029)* fixed two-cell embryos treated with the indicated RNAi and immunostained with PQN-59 antibodies (red). TIAR-1 GFP signal is in green and DNA was counterstained with DAPI (blue). Embryos were exposed to heat-shock (HS, 34°C for 10 minutes) before fixation. (**B**) Quantification of the average TIAR-1 granule number (left) and the average normalized TIAR-1 granule intensity (right) per embryo. Between 22 and 29 embryos were analyzed for each condition and genotype, N=3. Error bars indicate S.D. The P-value was determined using 2way ANOVA test. (**C**) Single confocal planes of *pqn-59::GFP;gtbp-1::RFP* fixed two-cell embryos treated with the indicated RNAi. DNA was counterstained with DAPI (blue). Embryos were exposed to 34°C for 10 mins (HS) or left at 20°C (no HS). For all images, scale bars represent 10 μm and enlarged ROIs are on the right. (**D**) and (**E**) Quantification of the average PQN-59 in (**D**) and GTBP-1 in (**E**) granule number (top) and the average normalized PQN-59 in (**D**) and GTBP-1 in (**E**) granule intensity (bottom) per embryo *(ctrl(RNAi)* n=22; *tiar-1(RNAi)* n=28, N=3). Error bars indicate S.D. The P-value was determined using Student’s t-test.

We then asked whether formation of PQN-59 and GTBP-1 granules is abolished when TIAR-1 is depleted. As shown in Fig 4C, the number of PQN-59 and GTBP-1 granules was not different between *tiar-1(RNAi)* and *ctrl(RNAi)* embryos exposed to heat-shock (quantifications in Figs 4D and 4E). The intensity of the signal of PQN-59 and GTBP-1 in the granules was weekly reduced (Figs 4D and 4E), a reduction that was not significant for GTBP-1::RFP. Immunostaining with αPQN-59 antibodies of embryos from the *tiar-1(tn1543)* mutant (17) exposed to heat-shock revealed a result similar to the RNAi depletion (S6B Fig).

These results indicate that TIAR-1 is not essential for assembly of PQN-59/GTBP-1 granules and that absence of both PQN-59 and GTBP-1, although associated with a reduced number of TIAR-1 granules, is not sufficient to abolish TIAR-1 granule assembly.

### PQN-59 is required for embryonic development and maintenance of brood size in a stress independent manner

PQN-59 and GTBP-1 both contribute to proper granule formation following heat-shock. We next investigated whether these two proteins are important for other functions in *C. elegans*, in normal growing conditions and therefore independently of a stress response.

We first asked whether brood size is reduced by the depletion of PQN-59 or GTBP-1. We found that depletion or null mutation of PQN-59 resulted in a significant reduction of progeny number whereas the depletion or null mutation of GTBP-1 did not (Figs 5A and 5B). Co-depleting both PQN-59 and GTBP-1 or depleting PQN-59 in the *gtbp-1(ax2029)* mutant resulted in a small but significant increase in brood size compared to the PQN-59 depletion alone (Fig 5A and S7A Fig).

**Fig 5.**
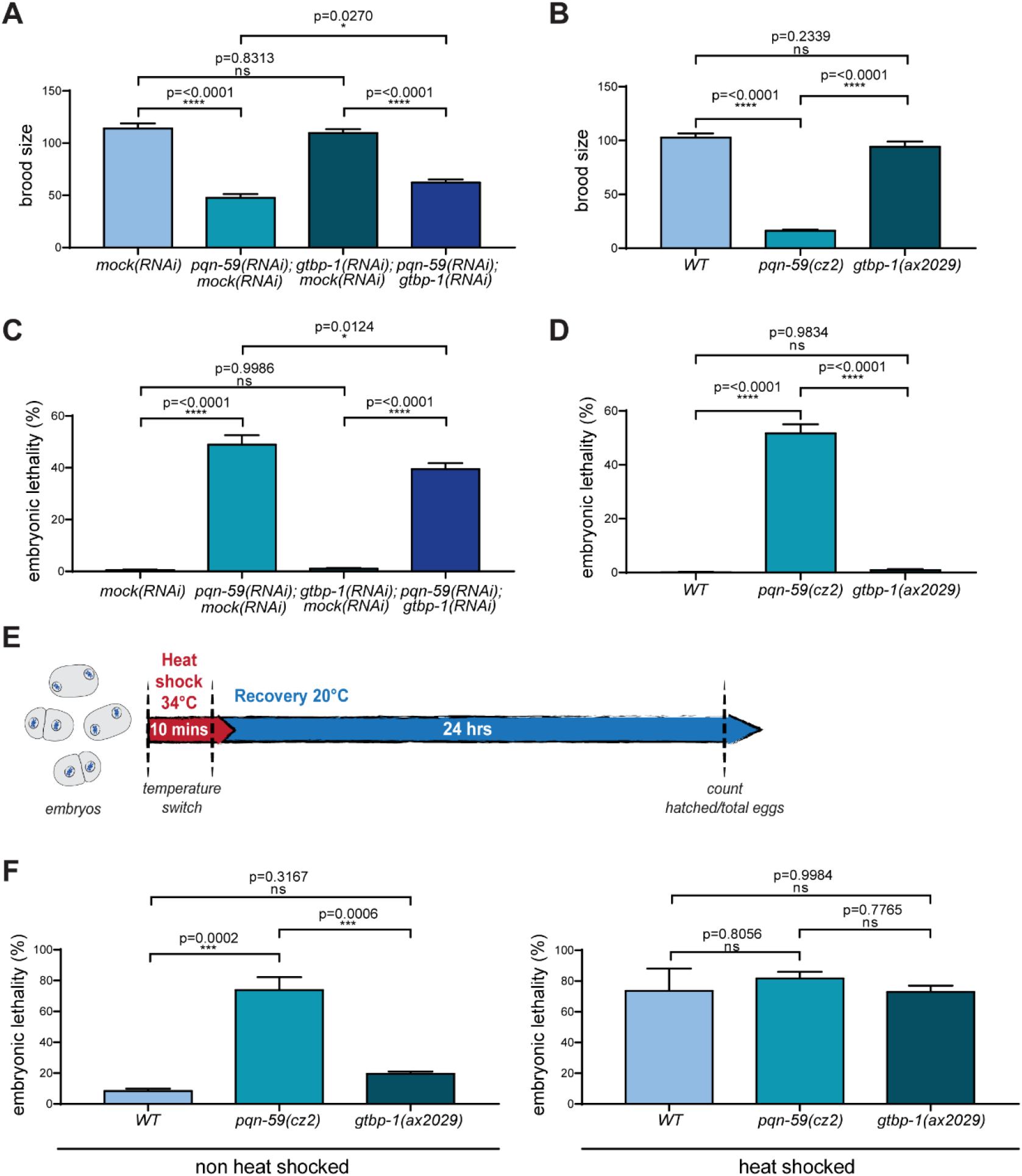
PQN-59 is important for C. elegans embryonic development and brood size. (**A**) and (**B**) Brood size of *wild-type* embryos after RNAi depletion of the indicated genes in (**A**) and of *wild-type,pqn-59(cz2)* and *gtbp-1(ax2029)* strains in (**B**). Values correspond to the average number of eggs laid per single worm. In **A** the brood size of a total of 30 animals per condition was evaluated in 3 independent experiments. In **B** the brood size of more than 30 animals of each genotype was evaluated in 2 or more independent experiments. (**C**) and (**D**) Embryonic lethality of *wild-type* embryos after RNAi depletion of the indicated genes in (**C**) and of *wild-type, pqn-59(cz2)* and *gtbp-1(ax2029)* strains in (**D**). Values correspond to the percentage of un-hatched embryos over the total progeny number (un-hatched embryos and larvae). In (**C)** embryonic lethality was assessed by counting more than 300 progeny for each condition, N=3. In (**D)** embryonic lethality was assessed by counting more than 200 progeny for each genotype, N>2. Error bars indicate S.E.M. The P-values were determined using one-way ANOVA test. (**E**) Timeline of the embryonic survival after heat-shock (see Materials and methods for more details). (**F**) Lethality of embryos exposed or not to heat shock was assessed as in (**E**) for *wild-type, pqn-59(cz2)* and *gtbp-1(ax2029)* strains counting the un-hatched embryos over the total number of embryos. The percentage of embryonic lethality is represented in **F**. More than 60 embryos per condition and genotype were counted in three independent experiments (N=3). Error bars indicate S.E.M. The P-values were determined using one-way ANOVA test.

We also found that depleting PQN-59 resulted in about 50% embryonic lethality (Fig 5C) a value similar to the PQN-59 mutant (Fig 5D). These results suggest that PQN-59 has an important function during embryonic development. On the contrary, depletion or mutation of GTBP-1 did not result in significant embryonic lethality (Figs 5C and 5D). Depleting GTBP-1 did not increase lethality of *pqn-59(RNAi)* embryos compared to the depletion of PQN-59 alone (Fig 5C), it actually weakly rescued (see discussion). This result was confirmed by the depletion of PQN-59 in the *gtbp-1(ax2029)* mutant (Fig S7B).

We then dissected wild-type and mutant hermaphrodites, exposed embryos to 34°C for 10 minutes and analysed how this treatment (Fig 5E) impacted on their viability. After 24 hours of recovery at 20°C, we found that embryonic lethality ranged from about 70% to 80% and we did not detect a significant difference between the wild-type, able to assemble proper stress granules, and the mutant embryos (Fig 5F).

Taken together our results suggest that PQN-59 and GTBP-1 do not help embryos to better resist to exposure to heat. Our results also indicate that PQN-59 has additional roles in adult life and during development that are independent of GTBP-1 and stress granule formation.

## Discussion

Here we have studied the function of two conserved proteins, PQN-59, the ortholog of UBAP2L, and GTBP-1, the ortholog of G3BP1/2 in assembly of stress granules in worm embryos and in worm germlines.

Both PQN-59 and GTBP-1 are cytoplasmic proteins that condense into granules in response to stress exposure. In *Drosophila melanogaster,* Lingerer/PQN-59 and Rasputin/GTBP-1 interact in Yeast Two Hybrid assays (28). In human cells, G3BP-1 and UBAP2L coimmunoprecipitate and mutations in UBAP2L that abolish the interaction with G3BP-1, are unable to rescue the stress granule assembly defect of UBAP2L depletion (12,25). *C. elegans* GTBP-1 was isolated in pull down of PQN-59 from embryos (unpublished), and we found that PQN-59 and GTBP-1 interact in a Yeast Two-Hybrid assay, supporting the hypothesis that PQN-59 and GTBP-1 are in a complex in *C. elegans.* In contrast with their human orthologs, the interaction of these two proteins or their presence is not essential for the formation of stress-induced granules, as revealed by looking at GTBP-1 or PQN-59 and TIAR-1. However, the assembly of stress-induced granules in the absence of one or the other is impaired, as in this condition the granules appear less numerous, less defined in their shape, and show a faster dissolution timing after stress relief. This suggests that the association between PQN-59 and GTBP-1 is not essential to assemble stress-induced granules but it is important to preserve stress granule properties.

Deletion of the RGG domain of UBAP2L results in the abolishment of all interactions with stress granule components and impairs stress granule assembly (12). Here we show that deletion of the RGG domain in PQN-59 does not result in defects in the number of stress granules, suggesting that this domain is dispensable for stress granule nucleation in the *C. elegans* embryo.

Single depletion of GTBP-1 and PQN-59 did not reduce the average number of TIAR-1 granules but granule number was highly variable, suggesting that PQN-59 and GTBP-1 do contribute to proper TIAR-1 granule formation. Consistent with a contribution, when PQN-59 was depleted in a *gtbp-1* mutant, the number of TIAR-1 granules was reduced. This indicates that PQN-59 and GTBP-1 are not strictly essential for TIAR-1 stress-induced granule assembly but they facilitate their formation. On the opposite, depletion of TIAR-1 did not result in a significant defect in the number of GTBP-1 and PQN-59 granules, suggesting that TIAR-1 may act downstream in the process of stress granule formation in *C. elegans* embryos.

Altogether, our data show that none of these proteins is required for stress induced granule assembly. So, whereas in cultured human cells G3BPs and UBAP2L are important to form stress granules in many stress conditions (9–12,25,32), in *C. elegans,* stress-induced granules can form in the absence of GTBP-1, PQN-59, and in the absence of both suggesting that either an essential nucleator of stress granules has still to be identified in this model or that the presence of disordered proteins is sufficient to assemble stress induced granules in worms. This is reminiscent of work in intestinal progenitor cells in *Drosophila* where canonical nucleators are not required for stress granule formation (33).

Depletion and mutation of PQN-59 result in additional phenotypes such as slow growth, reduced progeny and embryonic lethality, all in absence of stress. These phenotypes were not observed in *gtbp-1* mutant or depleted animals. A recent paper has shown that the human orthologs, G3BP1/2 inhibit mTORC1 signaling by targeting mTORC1 to the lysosome (34). One possibility is that the phenotypes of *pqn-59* mutant embryos are dependent on GTBP-1. For example, an excess of free GTBP-1 (not in complex with PQN-59) could be deleterious for worms and embryos. Co-depletion of both PQN-59 and GTBP-1 resulted in a weak rescue of the embryonic lethality and the reduced progeny phenotypes of *pqn-59* mutants, indicating that these phenotypes may partially depend on an excess of free GTBP-1. However, this weak rescue suggests that PQN-59 has additional important functions in embryos and worms that do not depend on GTBP-1. These yet to be identified functions could contribute to the regulation of the response to stress or be completely independent on the role of PQN-59 in stress granule assembly. Additional studies will be required to understand the molecular functions of PQN-59.

Stress granules have been proposed to protect cells from stress. We find that exposure to heat stress kills to the same extent wild-type, *pqn-59* or *gtbp-1* mutant embryos. This indicates that during embryonic development, the exposure to heat stress results in developmental failure, whether embryos are able to form proper stress granules or not.

## Materials and Methods

### Strains

The *C. elegans* strains used in this work are listed in Table 1. Worms were maintained on NGM plates seeded with OP50 bacteria, using standard methods (35). All the strains were grown at 20°C and incubated at 20°C after dsRNAs injections.

**Table 1.**
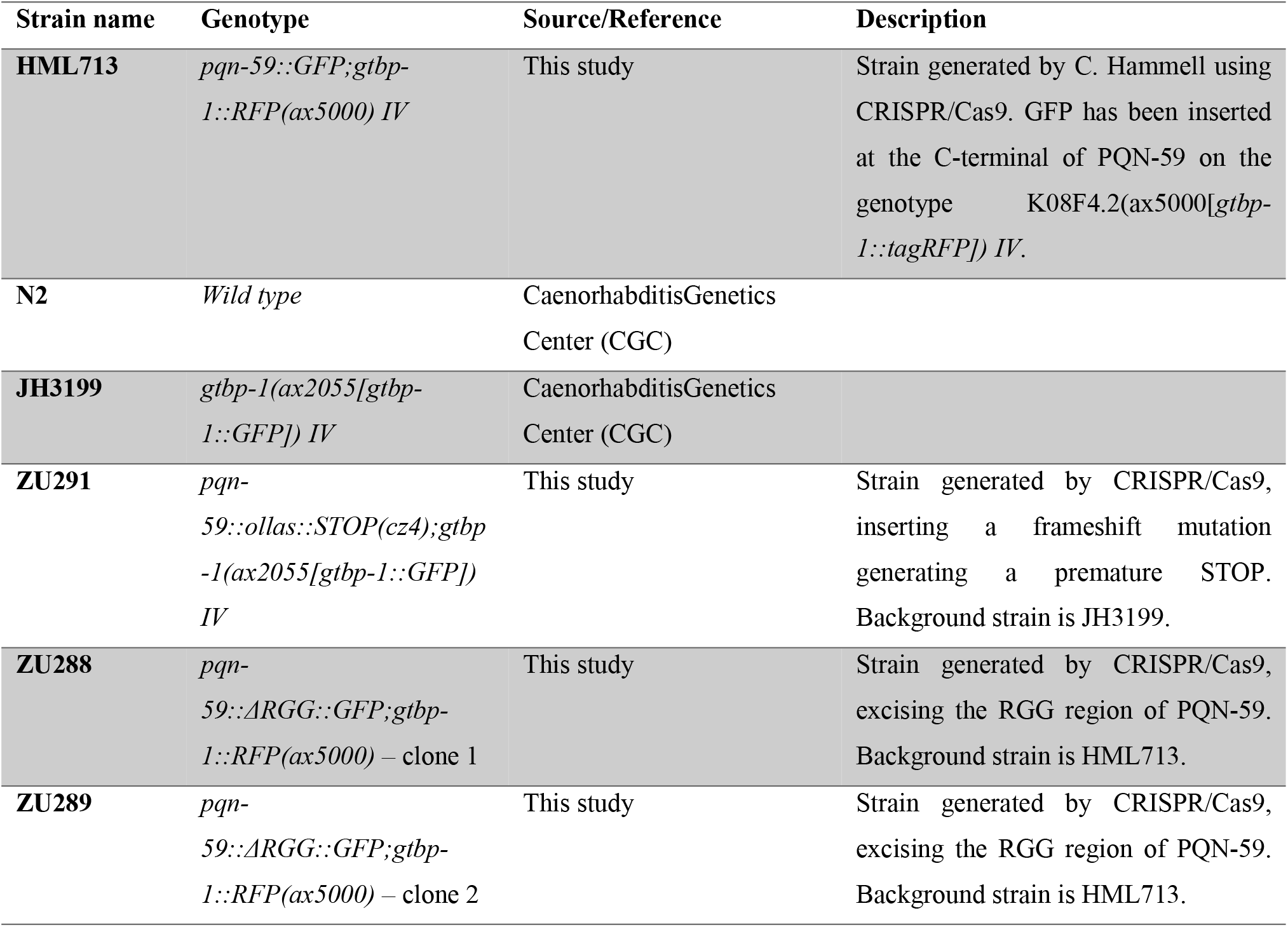
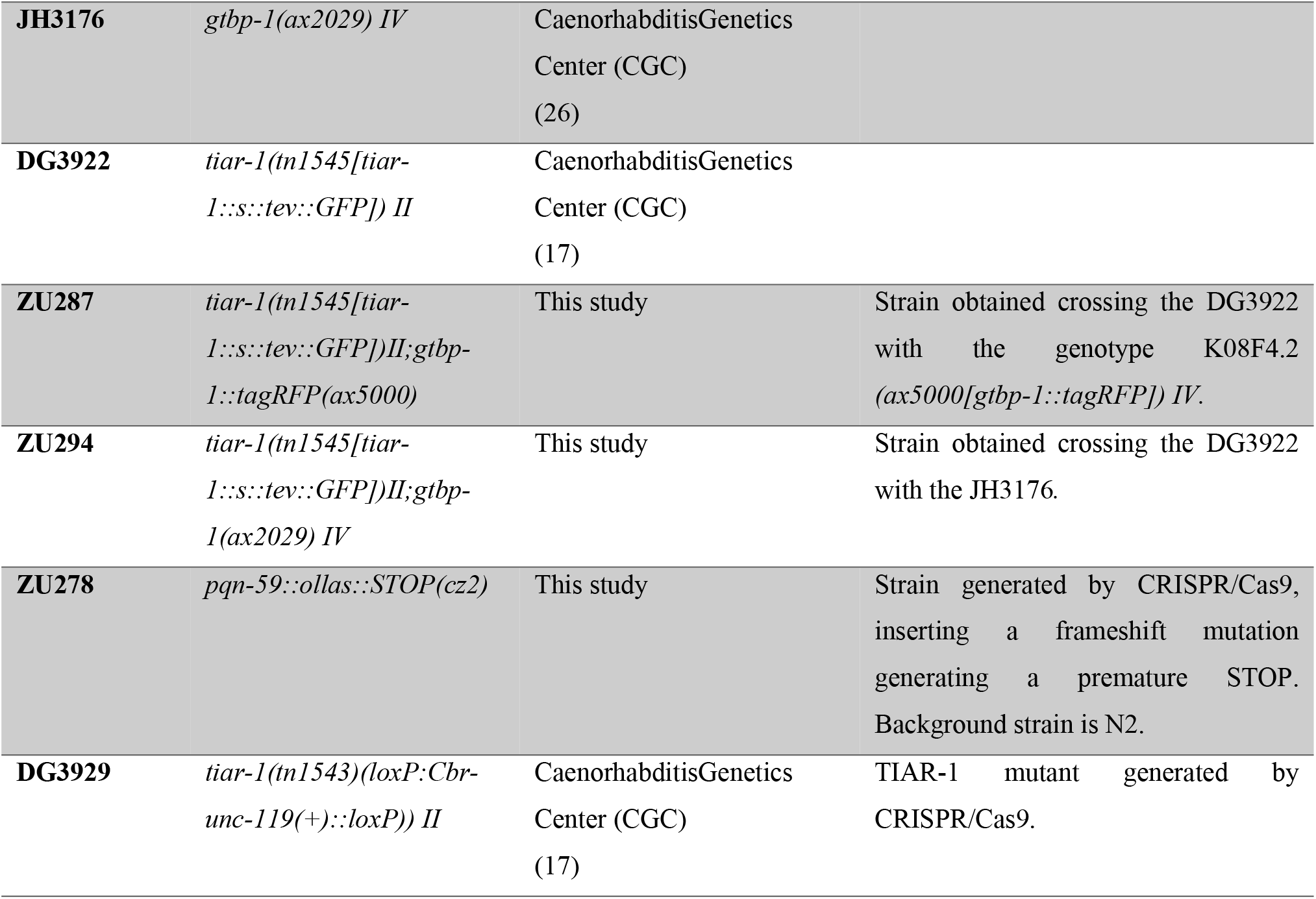
Strain List. The strain used in this work are listed in this table in order of appearance in the text. The genotype, source and description of the mutation are also reported.

Mutant strains were generated using CRISPR/Cas-9 technology, as described in (36). Single-guide RNAs and repair templates, as well as PCR primers used to detect and sequence the mutations, are listed in Table 2 and Table 3 respectively. The *pqn-59* mutant strain (generated in the N2 background and in the JH3199 *(gtbp-1(ax2055[gtbp-1::GFP])IV)* background) was generated by introducing a frameshift mutation leading to the appearance of a premature STOP codon. The *pqn-59ΔRGG* strain was generated excising the RGG-rich region (from aminoacid position 122 to aa 189), not altering the reading frame.

**Table 2.**
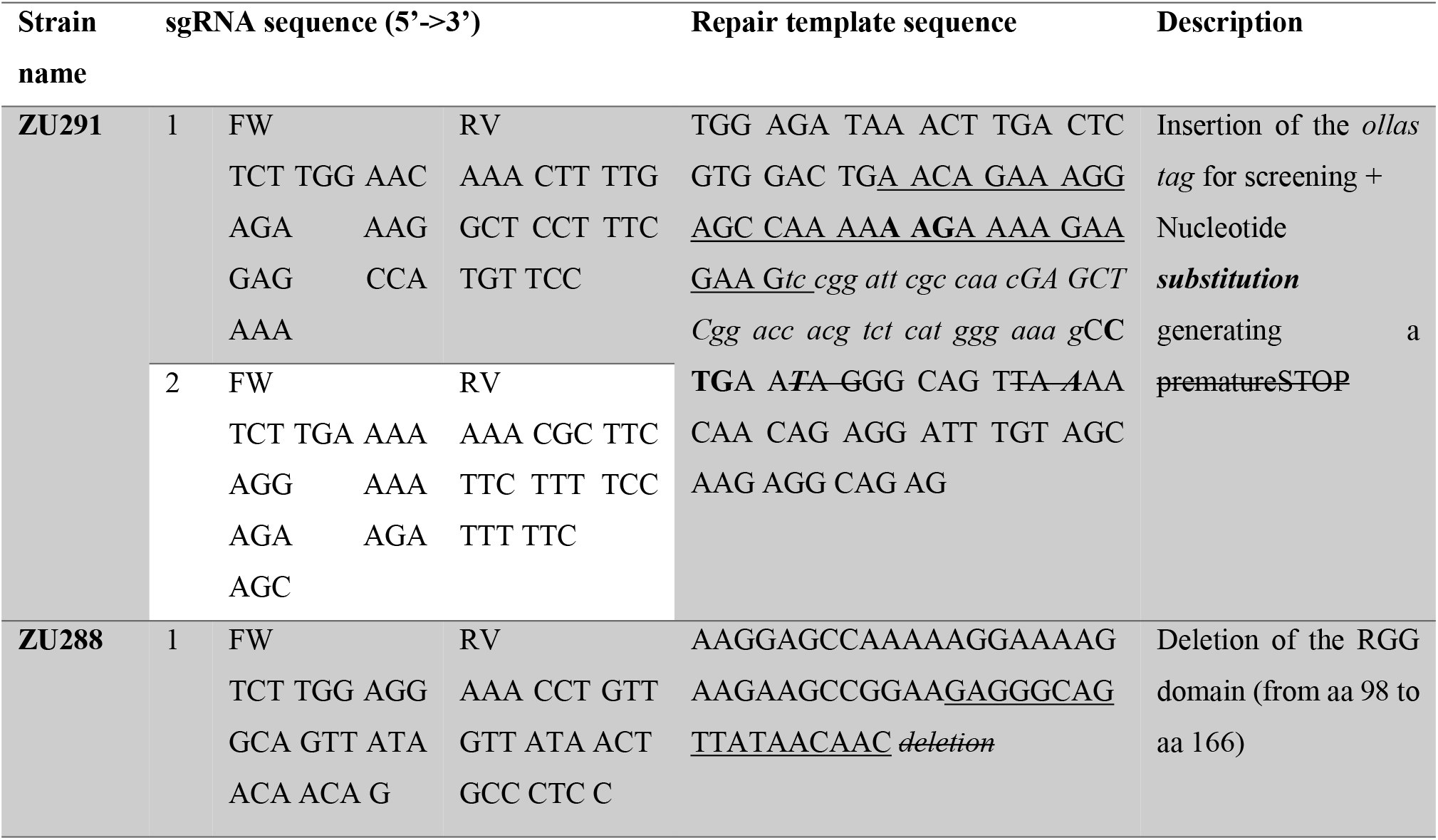
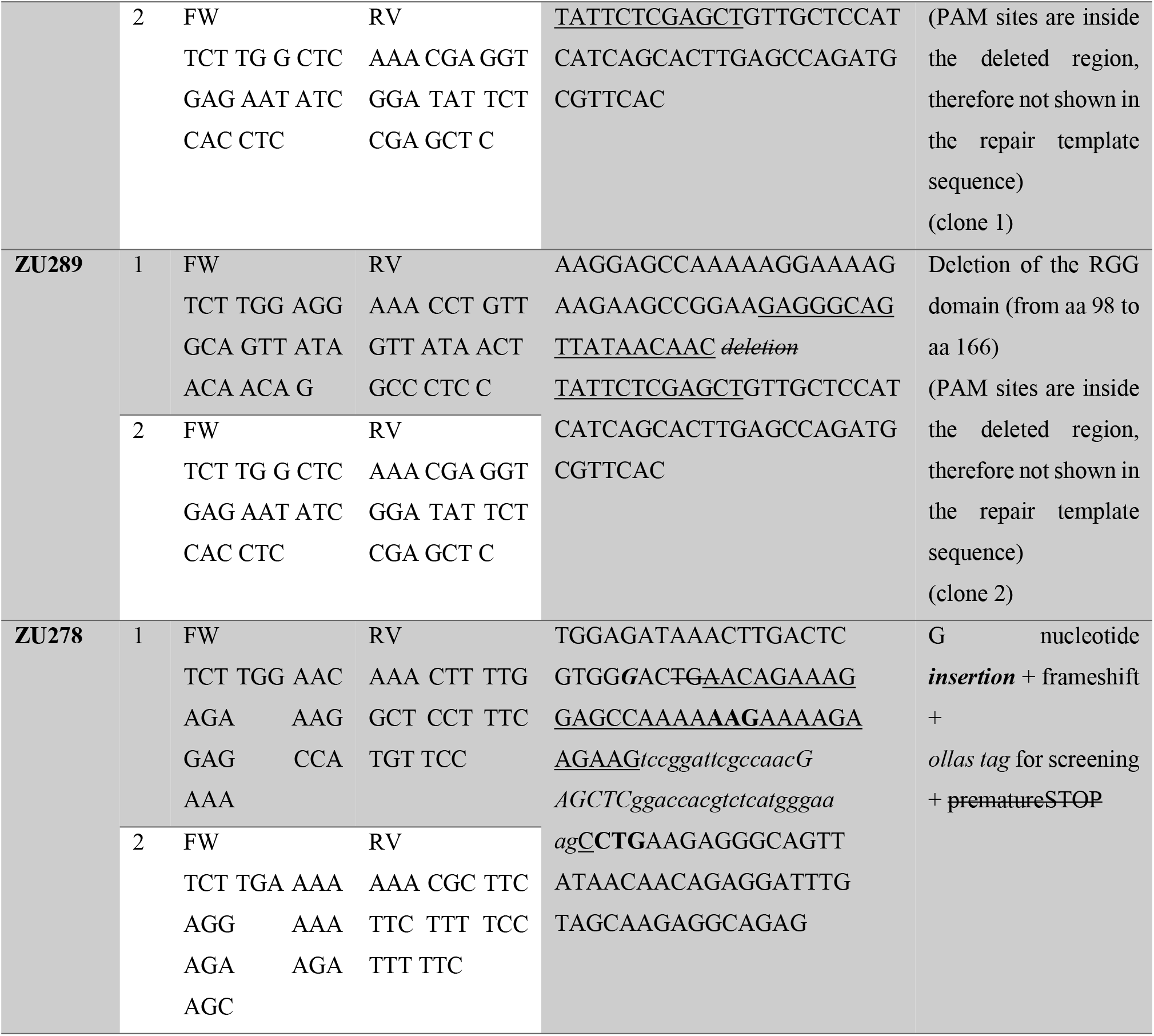
CRISPR reagents.

**Table 3.**
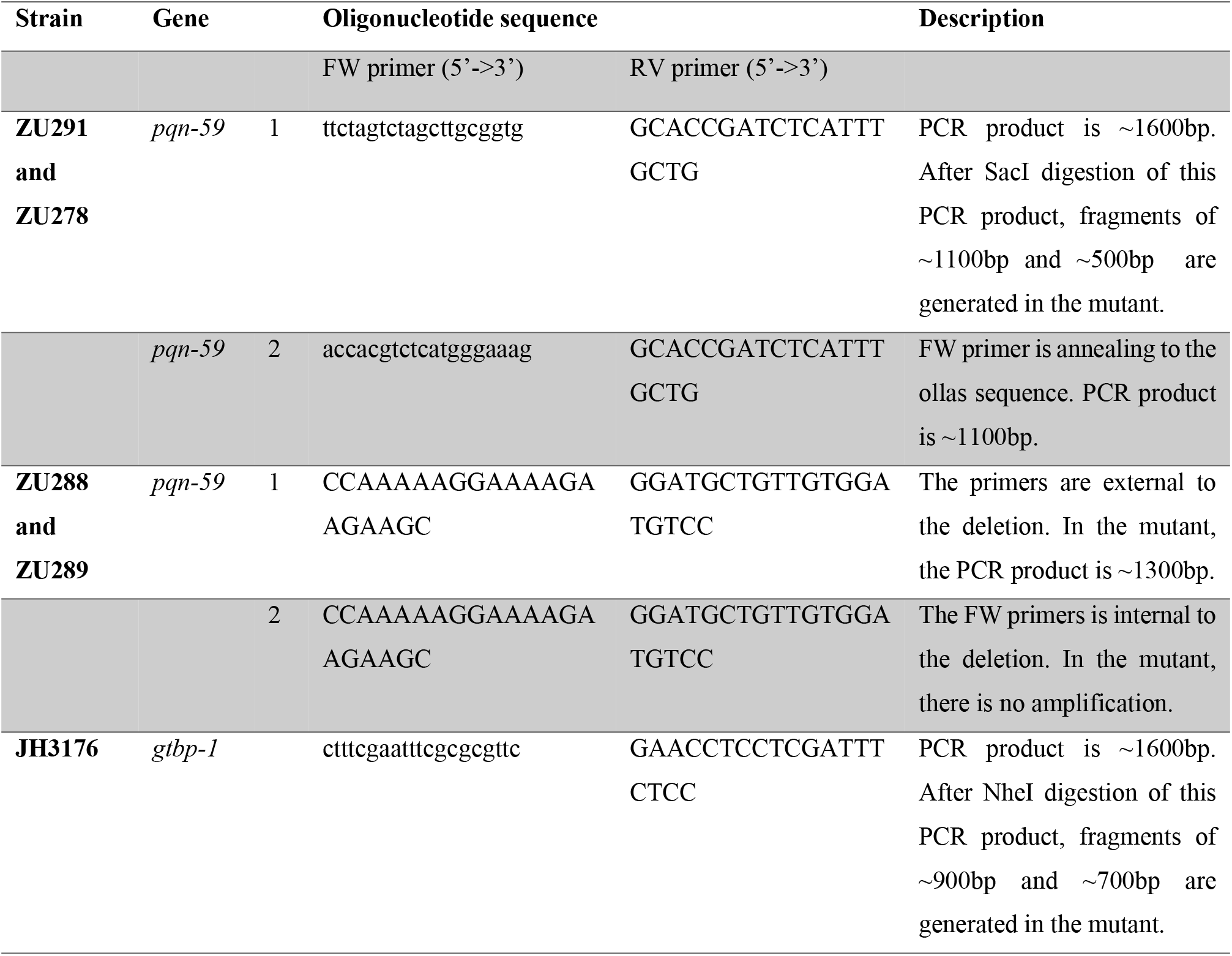
Oligonucleotides for PCR genotyping and sequencing.

### LABEL

<u>Underlined sequence</u> is targeted by the sgRNA

In capital bold the silently mutated **PAM sites** (the first one originally AGG mutated in AAG and the second one originally CGG mutated in CTG)

in italic is the *OLLAS* sequence containing a SacI restriction site (italic upper cases)

In bold capital italic the ***nucleotide substitution*** or ***insertion***

Strikethrough is the premature STOP codon and the deletion

### RNA interference

A list of the genes silenced through RNAi in this study is provided in Table 4.

**Table 4.**
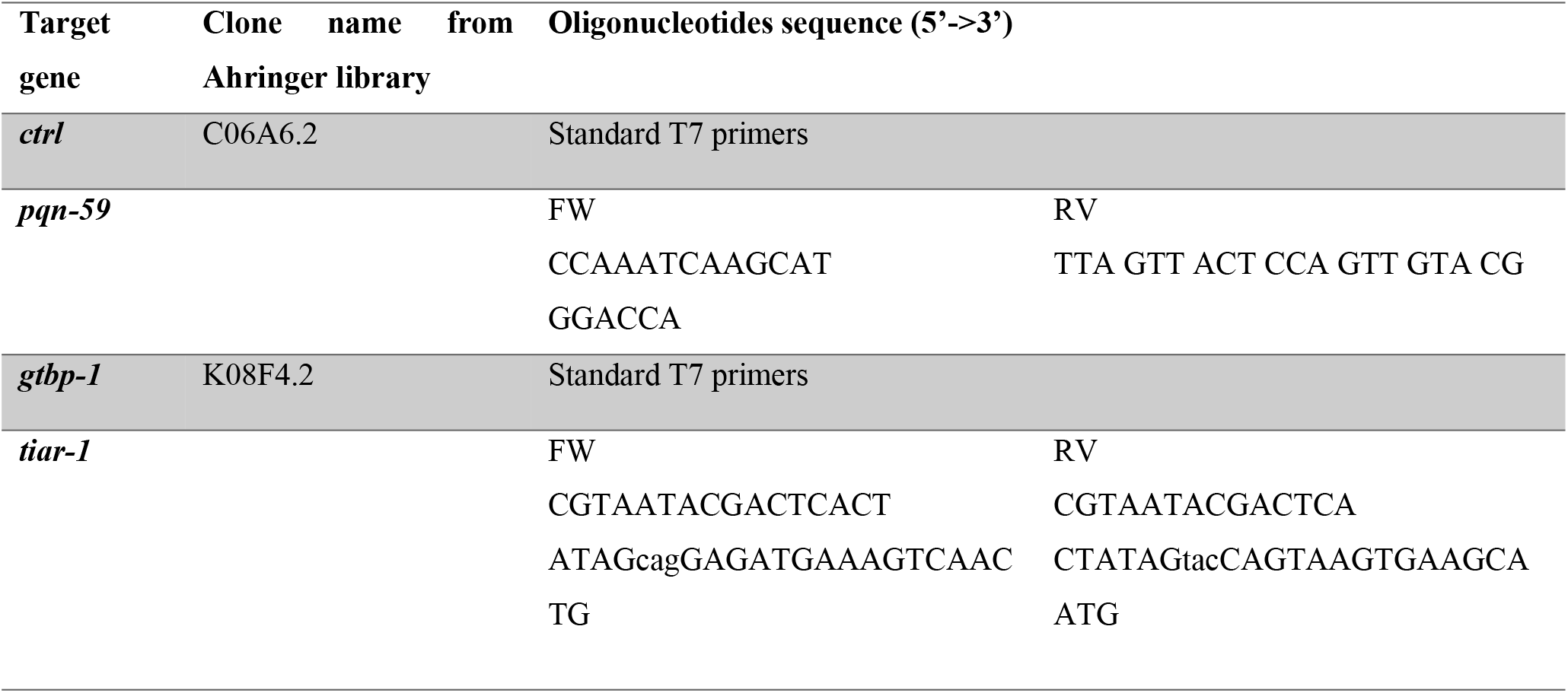
dsRNA sequences.

Clones from the Ahringer feeding library (37,38) were used when available. As a control, we used the clone C06A6.2, previously found in the laboratory to not affect early embryonic division and development (injected worms are 100% viable). To produce *pqn-59* dsRNA, a DNA fragment was amplified from genomic DNA using Gateway-compatible oligonucleotide primers (as in Table 4) for Gateway-based-cloning into the pDESTL4440 plasmid. The DNA was subsequently amplified using standard T7 primers. For *tiar-1*, the DNA was amplified from genomic DNA using oligos with T7 overhangs (see Table 4). For all genes, the dsRNA was produced with the Promega Ribomax RNA production system. dsRNA was injected in L4/young adult hermaphrodites which were incubated at 20°C. Germlines or embryos collected from injected hermaphrodites were analyzed 24 hours after injection.

### Live imaging of embryos exposed to heat-shock

Gravid hermaphrodites were dissected on a coverslip into a drop of Egg Buffer (118 mM NaCl, 48 mM KCl, 2 mM CaCl_2_, 2 mM MgCl_2_, and 25 mM Hepes, pH 7.5) containing 1:10 volume of polystyrene beads (Polybead® Hollow Microspheres, Polysciences). The temperature controller CherryTemp (Cherry Biotech, Rennes, France) with its accompanying software (Cherry Biotech TC) was used to control the temperature during the live imaging process. The coverslip with dissected hermaphrodites was directly mounted on the chip of the CherryTemp microfluidic temperature control system. The system was mounted on a Leica DM6000 microscope, equipped with epifluorescence and DIC (Differential Interference Contrast) optics and a DFC 360 FX camera (Leica). Time lapse images were collected every 10 seconds using 63x/1.4 numerical aperture (NA) objective and LAS AF software (Leica Biosystems). Imaging was started at 20°C. The temperature was then shifted at 30°C (heat-shock) for 5 to 10 minutes while imaging. For recovery, the temperature was shifted back to 20°C for 15-20 minutes.

### Immunostaining of embryos and image acquisition

For *C. elegans* embryos staining, 20-25 gravid hermaphrodites were dissected in a drop of M9 (86 mM NaCl, 42 mM Na2HPO4, 22 mM KH2PO4, and 1 mM MgSO4) on 22 × 40 mm coverslips.

Control samples were left at room temperature (20-22°C) for 10 minutes. For heat-shock exposure, coverslips with dissected worms and embryos, were transferred on a metal block placed in a humidified incubator at 34°C for 10 minutes.

After the incubation time, the coverslip was mounted crosswise on the epoxy slide square, previously coated with 0.1% poly-L-lysine, for embryonic squashing. The slides were then transferred on a metal block on dry ice for at least 10 min. Afterward, the coverslip was removed (freeze-cracking method) before fixation. Immunostaining was performed as described in (39). Briefly, embryos were fixed for 20 min in methanol and placed for 20 min in a solution of PBS and 0.2% Tween (PBST) and BSA 1% to block the nonspecific antibody binding. The slides were incubated with primary antibodies diluted in PBST with 1% BSA overnight at 4°C. The list of primary antibodies used in this study is in Table 5. After two washes of 10 min each in PBST, slides were incubated for 45 min at 37°C with a solution containing secondary antibodies (4 μ/ml Alexa Fluor 488– and/or 568–coupled anti-rabbit or anti-mouse antibodies from Molecular Probes) and 1 μg/ml DAPI to visualize DNA in PBST. Slides were then washed two times for 10 min in PBST before mounting using Mowiol (30% wt/vol glycerol, 3.87 mM Mowiol [Calbiochem, 475904], 0.2 M Tris, pH 8.5, and 0.1% DABCO).

**Table 5.**
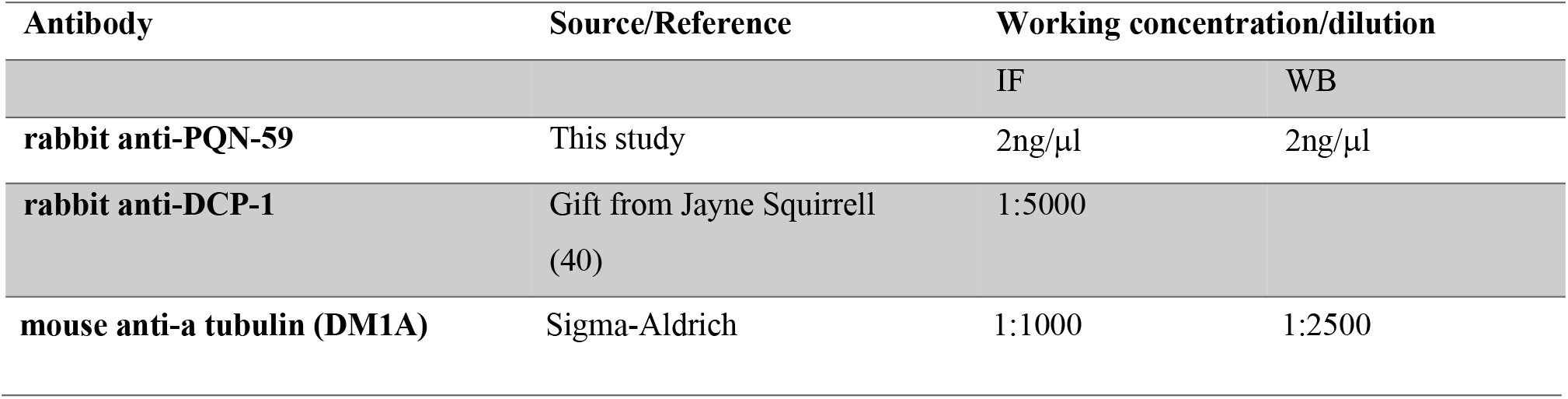
Primary antibodies list.

In the case of GFP or RFP tagged strains, the slides were briefly (10 min at room temperature) incubated with 1 μg/ml DAPI in PBST to visualize DNA just after methanol fixation and blocking. Slides were then washed two times for 10 min in PBST and mounted with Mowiol.

Images were acquired using a Nikon A1r spectral (inverted Ti Eclipse) confocal microscope equipped with a 60×1.4 NA CFI Plan Apochromat Lambda oil objective and four PMTS including two highly sensitive detectors (GaAsp) for green and red channels. From 5 to 7 z stacks, separated by 0.5 μm, were acquired. NIS Elements AR software (v.4.20.01; Nikon) was used to set acquisition parameters.

### Hermaphrodite heat-shock, drug treatment, and image acquisition procedure

For heat-shock, young adult worms were transferred into a drop of M9 buffer (86 mM NaCl, 42 mM Na_2_HPO_4_, 22 mM KH_2_PO_4_, and 1 mM MgSO_4_) on a glass coverslip and transferred on a metal block placed into a humidified incubator for 10 minutes at 35°C. For recovery after heat-shock, worms were collected from the M9 drop and transferred onto OP50 seeded NGM plates and incubated at 20°C for 5 or 10 minutes.

For drug treatment, young adult worms were transferred into a drop of M9 buffer only (control) or M9 with 10 mg/ml of Puromycin (InvivoGen) or with 20 mM Arsenite (MerckMillipore). Worms were incubated in the Puromycin-containing solution for 4 hours and in the Arsenite-containing solution for 5 hours before imaging. Control worms were incubated in M9 buffer for the same amount of time as the Puromycin or Arsenite treated worms.

Control and drug-treated worms were then transferred in a drop of NaN_3_ 30 mM (for worm paralysis) and mounted on a 3% agarose pad for imaging. Imaging was performed using the Leica DM6000 described above. Images were acquired using the 63x/1.4 numerical aperture (NA) objective and the LAS AF software (Leica Biosystems).

### Quantification of cytoplasmic protein levels

The mean intensity of a defined region of interest (ROI) (w=2.69, h=2.46, area=6.604), always placed in the anterior blastomere (AB) of a two-cell stage *C. elegans* embryo, was measured using Fiji Image J. The mean intensity of an equal ROI, placed outward of the embryo, was used for background subtraction. For each experiment, the obtained mean intensity values were normalized on the highest value for 0 to 100 (%) scale conversion.

### Quantification of cytoplasmic granules

For the quantification of PQN-59, GTBP-1 and TIAR-1 cytoplasmic granules QuPath version 0.2.3 was used (41). The algorithm for granule detection was based on a pixel classifier and was trained on representative pictures with dedicated annotations. For each embryo, manually delineated, the total number of detected granules was obtained. The average intensity of all the detected granules in each embryo was background subtracted using the average embryonic intensity of the same embryo.

### Yeast two-hybrid assay

The interaction between PQN-59 and GTBP-1 was assessed in the PJ69-4a yeast strain (42) using single copy GAL4-activiation and GAL4-DNA-binding domain-based vectors. Full-length cDNAs were cloned into these vectors using Gibson reactions and transformed into the host yeast strain using previously described protocols (42). Transformants were selected on SC-leu-trp plates and subsequently tested for growth (3 days) on SC-trp-leu-his plates containing 3mM 3AT.

### Protein domain identification

Protein domains were identified using the meta site Motif Scan tool, a free database for protein motif prediction developed by the Swiss Institute of Bioinformatics (SIB), including Prosite, Pfam, and HAMAP profiles (https://myhits.isb-sib.ch/cgi-bin/motif_scan). Comparable results have also been obtained interrogating other online tools, such as PROSITE at ExPASy (https://prosite.expasy.org/), MOTIF (GenomeNet, Institute for Chemical Research, Kyoto University, Japan) (https://www.genome.jp/tools/motif/), and InterPro (http://www.ebi.ac.uk/interpro/).

Prion domains have been identified using PLAAC (http://plaac.wi.mit.edu).

### Antibody production

To produce antibodies to PQN-59 a C-terminal fragment (aminoacid 304-712) was cloned using the Gateway technology (Invitrogen) into the pDEST15. Recombinant GST-tagged PQN-59 was expressed in BL21 and purified using standard protocols. Antibody production in rabbit was performed by Covalab, France. The obtained anti-PQN-59 serum was purified on membrane strip carrying bacterially expressed GST-PQN-59 antigen. About 5 μg of fusion protein was loaded in each lane of a 10% acrylamide gel. The protein was transferred on a nitrocellulose membrane (GE Healthcare). A stripe of the membrane, containing the protein, was cut and incubated for 1 hour in PBS + 3% milk for blocking. The band was then incubated overnight at 4°C in 1 ml of serum diluted in 1 ml of 3% milk in PBS-Tween + 4 mg of GST (to avoid GST binding). After three washes of 5 to 10 minutes, the antibody was eluted using a solution of glycine 100 mM, pH 2. The pH of the elution solution was equilibrated to 7.5 using TRIS 1M.

### Western Blot

For western blot, 50 adult worms were manually picked from NGM plates, resuspended in Laemmli sample buffer and denatured at 92°C for 2 minutes. Lysates were separated by SDS-PAGE using a 10% Acrylamide Gel. Proteins were then transferred onto a nitrocellulose membrane (Sigma). The membrane was blocked with 3% milk in PBS. After washing with a solution of PBS and 0.1% Tween (PBST), the membrane was incubated overnight at 4°C with primary antibodies diluted in a 1% BSA-PBST solution. The following day the membrane was washed with PBST twice for 10 minutes and incubated with secondary antibodies diluted in the same solution (1:10000 HRP-conjugated anti-mouse or anti-rabbit antibodies (Biorad)) in the same solution at room temperature for 45 minutes. After three washes of 10 minutes each, proteins were visualized with ECL (Millipore) using a Pxie machine.

### Embryonic lethality and brood size counting

To count embryonic lethality and brood size, L4/young adult worms were singled onto individual OP50-seeded NGM plates and incubated at 20°C for 24 hours. After 24 hours, the adult worm was removed, and plates were again incubated at 20°C for 24 hours. To assess the brood size, the total number of un-hatched embryos and hatched larvae were counted under a dissecting microscope. The ratio between the un-hatched embryos over the total of the F1 progeny (brood size) was used to calculate the percentage of embryonic lethality.

### Embryonic lethality after heat-shock

For embryonic lethality after heat-shock, gravid hermaphrodites were dissected on a coverslip into a drop of Egg Buffer (118 mM NaCl, 48 mM KCl, 2 mM CaCl_2_, 2 mM MgCl_2_, and 25 mM Hepes, pH 7.5) where the embryos were released. The coverslip was then transferred on a metal block placed in a humidified incubator at 34°C for 10 minutes. After the heat-shock, the embryos were transferred by pipetting on OP50-seeded NGM plates, counted, and incubated at 20°C for 24 hours for recovery. After recovery, the number of un-hatched embryos was counted. The ratio between un-hatched embryos over the number of embryos plated was used to calculate the percentage of embryonic lethality after heat-shock exposure.

### Statistical analysis

Statistical analysis was performed using GraphPad Prism 8. Details on the statistical test, the sample, and experiment number, as well as the meaning of error bars, are provided for each experiment in the corresponding figure legend, in the results and/or in the method details. Significance was defined as, ns, p > 0.05, *p < 0.05, **p < 0.01, ***p < 0.001, ****p < 0.0001.

## Acknowledgements

We would like to thank G. Seydoux (Johns Hopkins University) and Jayne Squirrell (University of Winsconsin) for strains and reagents. We thank present and past members of the Gotta laboratory for help, discussions and comments on the manuscript, with special thanks to Luca Cirillo (Institute of Cancer Research, ICR). We thank Patrick Meraldi, Florian Steiner and their laboratories for interesting discussions, suggestions and comments on the manuscript. Thanks to the Bioimaging Facility of the Medical Faculty and special thanks to Nicolas Liaudet for help with quantifications of stress granules. Some strains were provided by the CGC, which is funded by the NIH office of research infrastructure program (P40OD010440).

## Supporting information captions

**S1 Fig.**
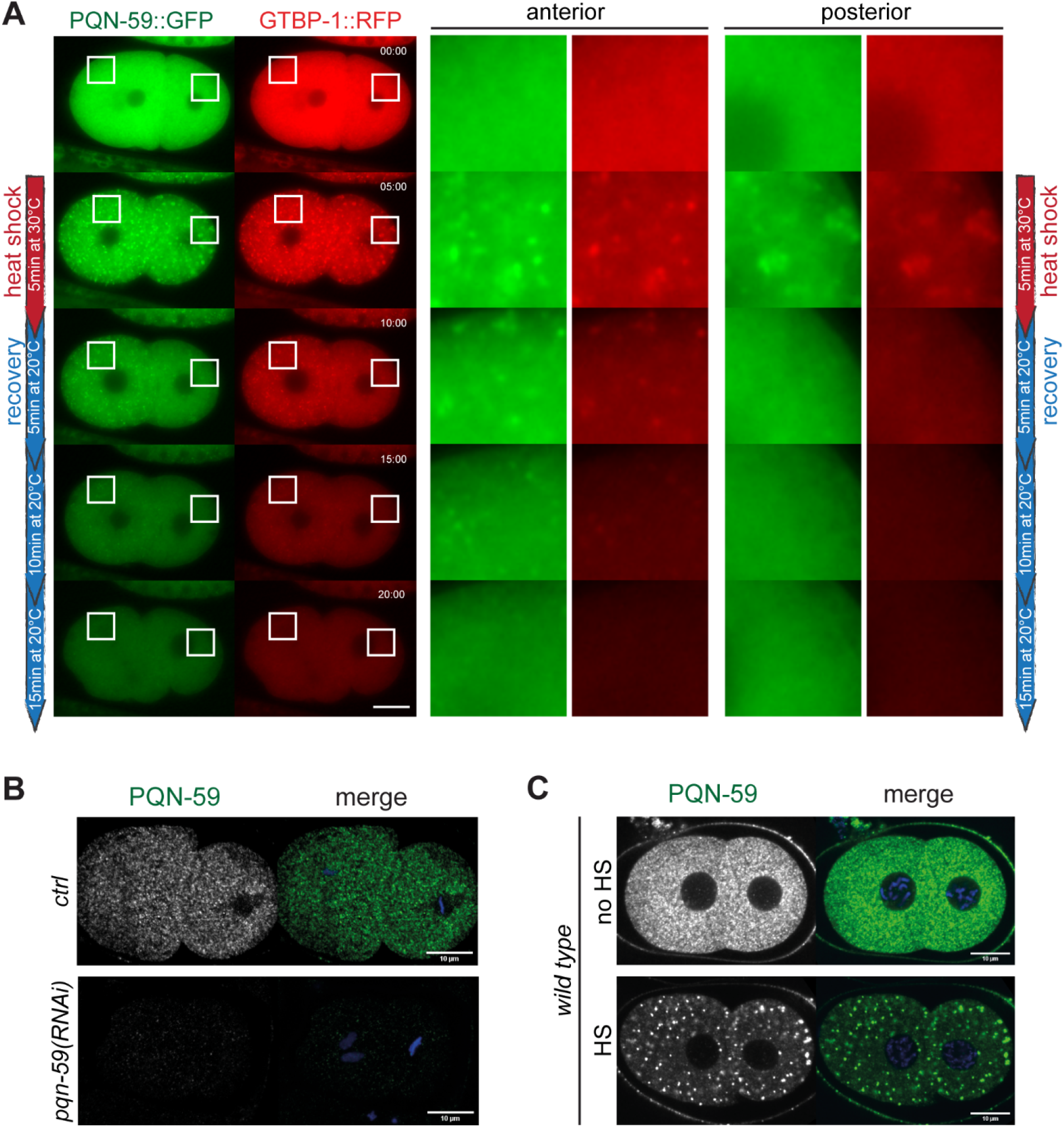
PQN-59/GTBP-1 heat-induced embryonic granules are reversible. (**A**) Still frames from time-lapse imaging of *pqn-59::GFP;gtbp-1::RFP* embryos using the CherryTemp temperature-controlled stage (n=12, N=3). The red vertical line on the left shows the time of exposure to HS and the blue line the time after stress release (recovery). (**B**) and (**C**) Fixed two-cell embryos immunostained with anti-PQN-59 antibodies (green). DNA was counterstained with DAPI (blue). (**B**) Maximum projections of confocal images of untreated *(ctrl)* or PQN-59-depleted *(pqn-59(RNAi))* embryos, as indicated (n=25 *ctrl* and n=17 *pqn-59(RNAi)* embryos, N=2). (**C**) Single confocal planes of two-cell *wild-type* embryos. Embryos were left at 20°C (no HS) or exposed to 34°C for 10 mins (HS) before fixation. (N=6). Scale bars represent 10 μm. Enlarged ROIs are on the right.

**S2 Fig.**
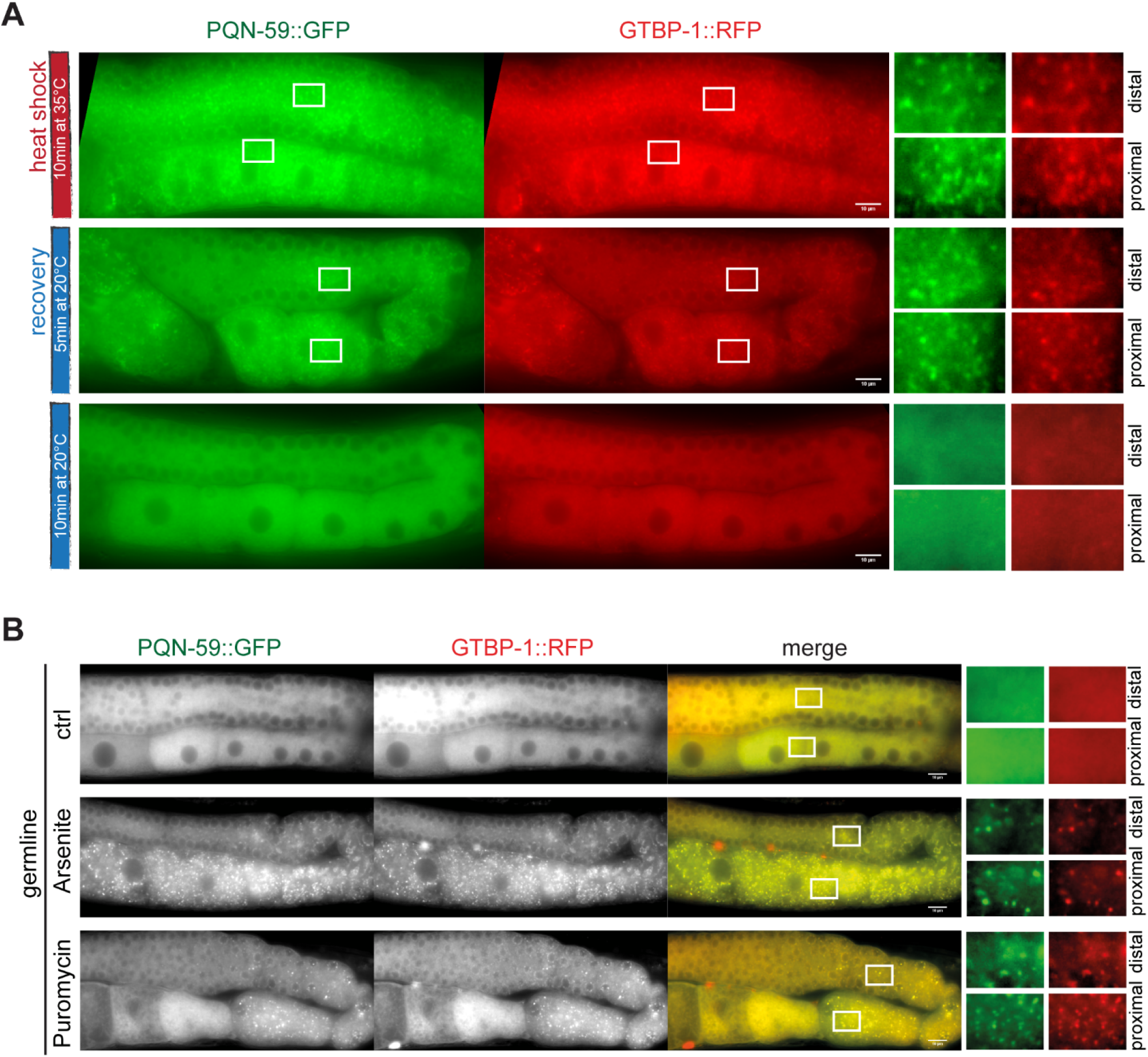
PQN-59/GTBP-1 granules form in response to several stresses in the germline and are reversible. (**A**) and (**B**) Images of germlines of *pqn-59::GFP;gtbp-1::RFP* adults. (**A**) PQN-59 (green) and GTBP-1 (red) form cytoplasmic granules after 5 minutes of heat exposure at 35°C (red vertical line) and dissolve after 10 minutes of recovery at 20°C (blue vertical line, n=15, N=3). (**B**) Control (buffer) and worms treated with drugs, as indicated on the left. 85% (n=26) of the Arsenite-treated worms and 100% of the Puromycin-treated worms (n=15) showed formation of PQN-59 (green) and GTBP-1 (red) granules in the germline. ROIs are shown enlarged on the right. Scale bars represent 10 μm.

**S3 Fig.**
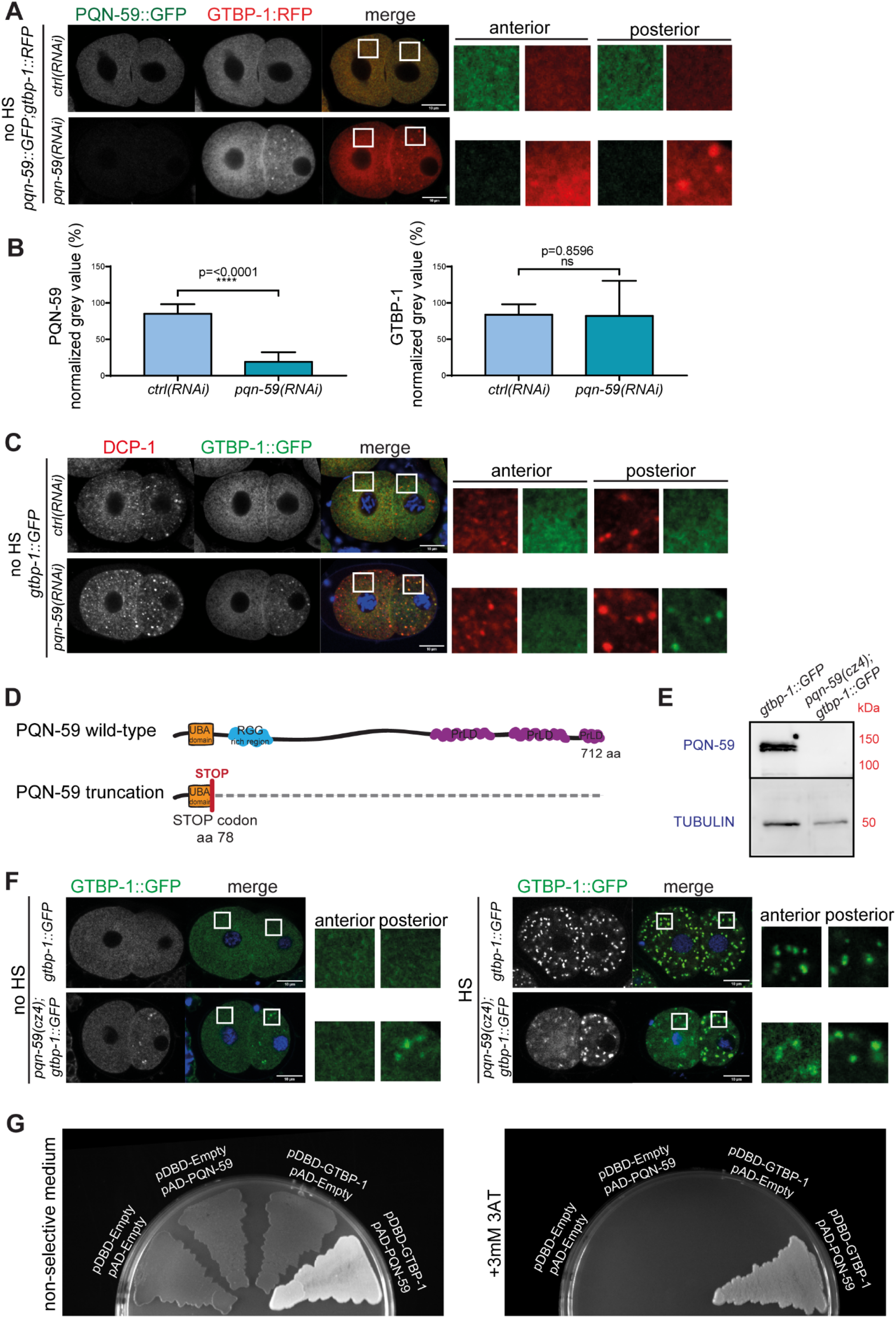
PQN-59 interacts with GTBP-1 and its depletion or deletion results in stress-independent GTBP-1 clusters in the posterior embryonic blastomere. (**A**) Single confocal planes of *pqn-59::GFP;gtbp-1::RFP* fixed two-cell stage embryos treated with the indicated RNAi in non-heat-shocked conditions (no HS). (**B**) Quantification of the normalized cytoplasmic intensity of PQN-59 (left) and GTBP-1 (right) of *ctrl(RNAi)* and *pqn-59(RNAi)* in the anterior blastomere as in (**A)** *(ctrl(RNAi)* n=23; *pqn-59(RNAi)* n=38, N=4). Error bars indicate S.D. The P-value was determined using Student’s t-test. (**C**) Single confocal planes of *gtbp-1::GFP* fixed two-cell stage embryos immunostained with DCP-1 antibodies (red) in non-heat-shocked conditions (no HS). GTBP-1 GFP signal is in green and DNA was counterstained with DAPI (blue). Embryos were treated with *ctrl* or *pqn-59(RNAi)* as indicated *(ctrl(RNAi)* n=12; *pqn-59(RNAi)* n=19, N=3). (**D**) Illustration of the PQN-59 protein domains with the indication of the STOP codon insertion in the strain *pqn-59(cz4);gtbp-1::GFP* obtained by CRISPR/Cas9. (**E**) Western blot on worm lysate of *gtbp-1::GFP* and *pqn-59(cz4);gtbp-1::GFP* worms using PQN-59 and TUBULIN (loading control) antibodies. (**F**) Single confocal planes of fixed two-cell stage *gtbp-1::GFP* and *pqn-59(cz4);gtbp-1::GFP* embryos, at 20°C (no HS) or after exposure to 34°C for 10 mins (HS). GTBP-1 GFP signal is in green and DNA was counterstained with DAPI (blue). ROIs are shown enlarged on the right of each set of embryos. Between 6 and 17 two-cell embryos were analyzed in each condition, N=4. Scale bars represent 10 μm. (**G**) Yeast two-hybrid assay using the PJ69-4a yeast strain transformed with the indicated plasmids. On non-selective plates, all streaks grow. Controls are in red because of lack of interaction and lack of activation of the ADE-2 reporter. The streak of cells containing both PQN-59 and GTBP-1 is white, indicating interaction-dependent activation of the ADE-2 reporter. On selective plates (+3mM 3AT) yeast growth is observed only for the clone where both PQN-59 and GTBP-1 are expressed, indicating interaction and activation of the HIS-3 reporter.

**S4 Fig.**
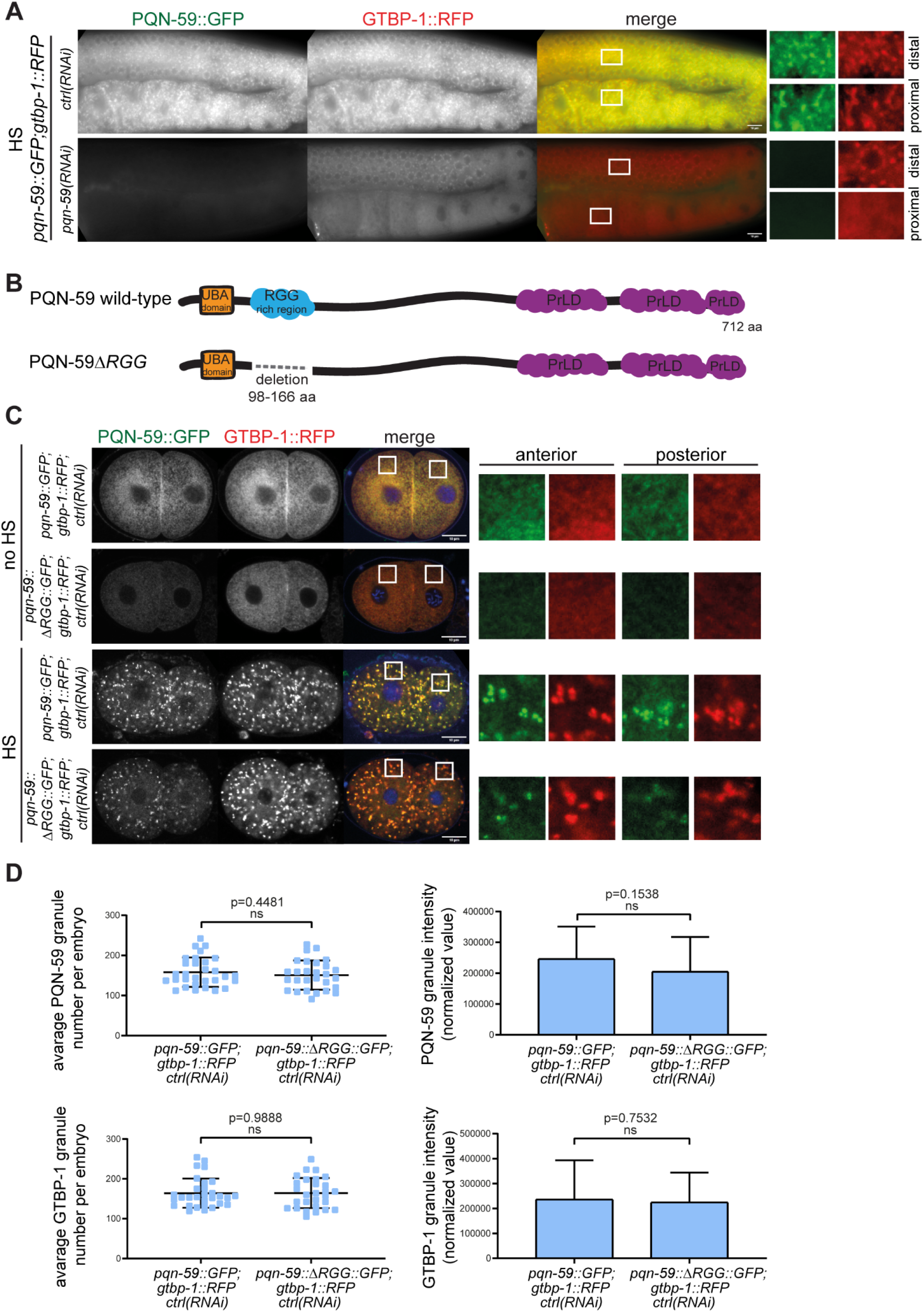
GTBP-1 stress-induced granule formation is impaired in the germline by depletion of PQN-59 but not in a strain in which the RGG domain has been deleted. (**A**) Germline pictures of *pqn-59::GFP;gtbp-1::RFP* worms treated with the indicated RNAi and exposed to heat-shock (HS, *ctrl(RNAi)* n=14; *pqn-59(RNAi)* n=15, N=2). ROIs are enlarged on the right. Scale bars represent 10 μm. (**B**) Illustration of the PQN-59 protein with the RGG deletion introduced by CRISPR/Cas9 in the strain *pqn-59::GFP;gtbp-1::RFP*. (**C**) Single confocal planes of *pqn-59::GFP;gtbp-1::RFP* and *pqn-59::ΔRGG::GFP;gtbp-1::RFP* fixed two-cell stage embryos. PQN-59 GFP signal is in green, GTBP-1 RFP signal is in red and DNA was counterstained in blue. On the right, ROIs are enlarged 8 X. Scale bars represent 10 μm. (**D**) Quantification of the average PQN-59 (top left) and GTBP-1 (bottom left) granule number per embryo and of the average normalized PQN-59 (top right) and GTBP-1 (bottom right) granule intensity per embryo *(pqn-59::GFP;gtbp-1::RFP* n=30; *pqn-59::ΔRGG::GFP;gtbp-1::RFP*, n=28, N=3). Error bars indicate S.D. The P-value was determined using Student’s t-test.

**S5 Fig.**
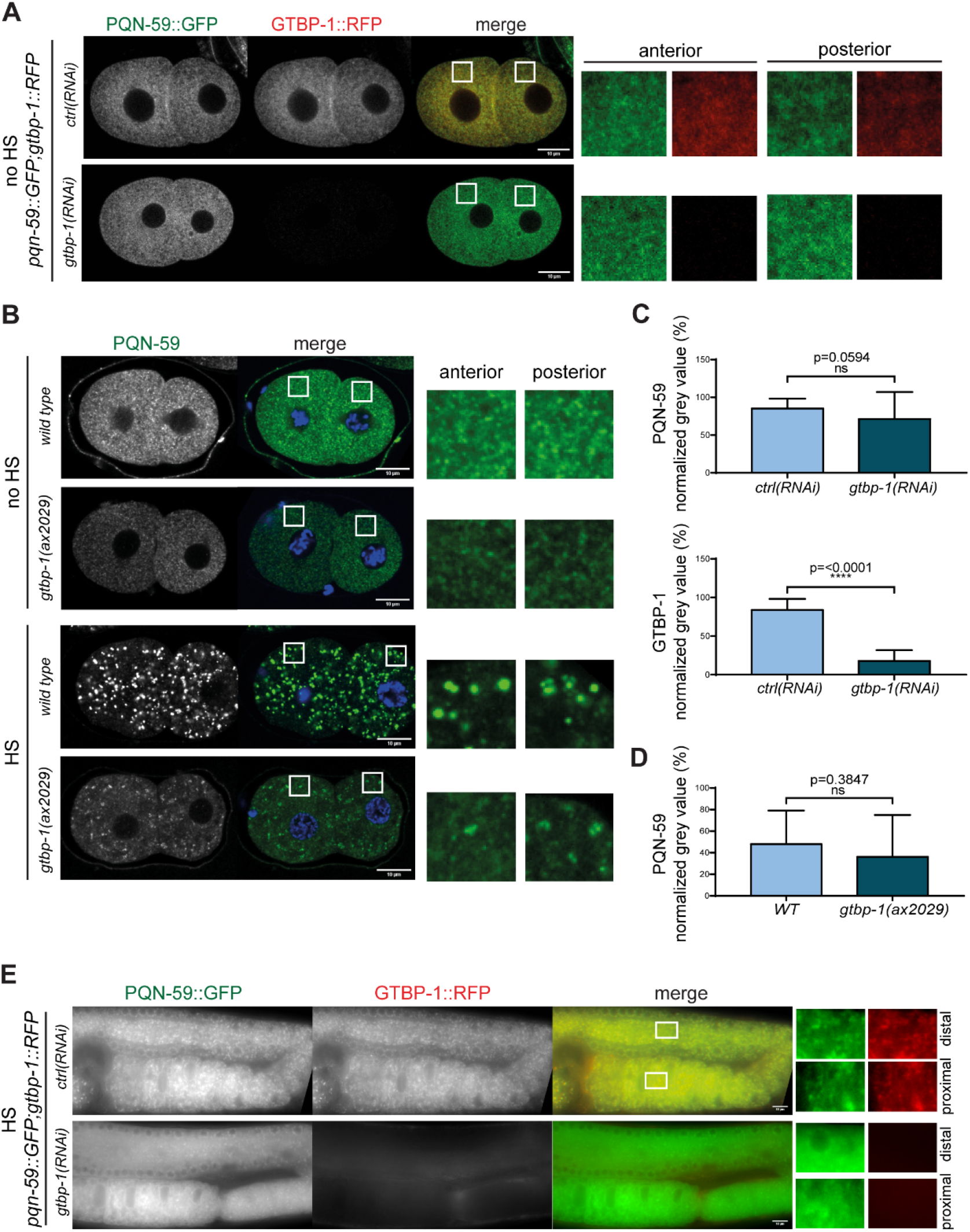
GTBP-1 null mutation impairs stress-induced PQN-59 granule formation. (**A**) Single confocal planes of *pqn-59::GFP;gtbp-1::RFP* fixed two-cell embryos treated with the indicated RNAi in non-heat-shocked conditions (no HS). (**B**) Single confocal planes of *wild-type* and *gtbp-1(ax2029)* fixed two-cell embryos immunostained with PQN-59 antibodies (green). DNA was counterstained with DAPI (blue). Embryos were exposed to 34°C for 10 mins (HS) or left at 20°C (no HS). For both (**A**) and (**B**) enlarged ROIs are shown on the right. Scale bars represent 10 μm. (**C**) Quantification of the normalized cytoplasmic intensity of PQN-59 (top) and GTBP-1 (bottom) of *ctrl(RNAi)* and *pqn-59(RNAi)* embryos at 20°C as in **A** *(ctrl(RNAi)* n=22; *gtbp-1(RNAi)* n=31, N=4). Error bars indicate S.D. The P-value was determined using Student’s t-test. (**D**) Quantification of the normalized cytoplasmic intensity of PQN-59 of *wild-type* and *gtbp-1(ax2029)* embryos at 20°C as in **B** *(wild type* n=14; *gtbp-1(ax2029)* n=14, N=3). Error bars indicate S.D. The P-value was determined using Student’s t-test. (**E**) Germline pictures of *pqn-59::GFP;gtbp-1::RFP* worms treated with the indicated RNAi and exposed to heat-shock (HS, *ctrl(RNAi)* n=15; *gtbp-1(RNAi)* n=16, N=2). ROIs are enlarged on the right.

**S6 Fig.**
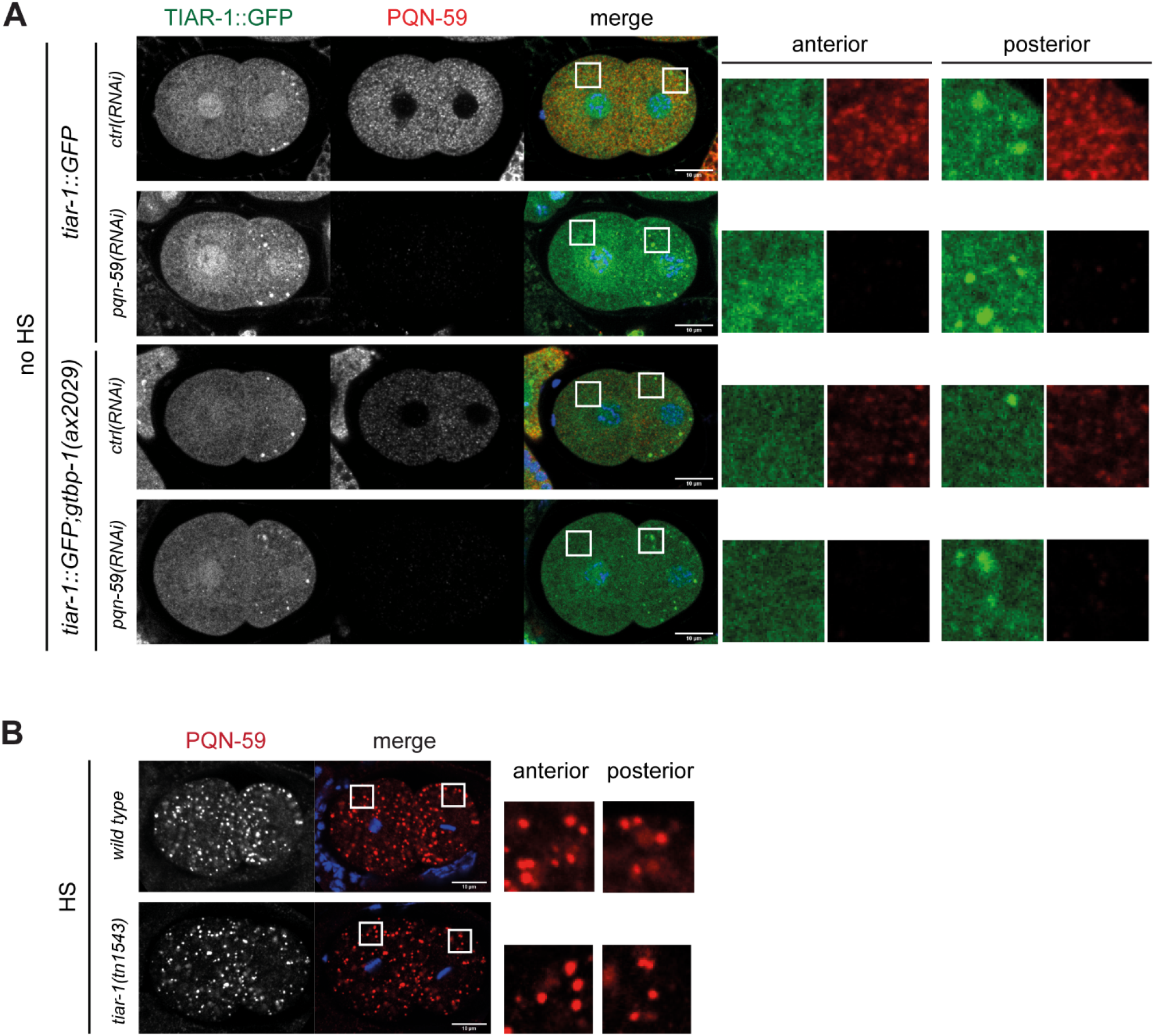
PQN-59 accumulation in stress-induced granules is not affected in *tiar-1* mutant embryos. (**A**) Single confocal planes of *tiar-1::GFP* and *tiar-1::GFP;gtbp-1(ax2029)* fixed two-cell embryos treated with the indicated RNAi in non-heat-shocked conditions (no HS). Embryos were immunostained with PQN-59 antibodies (red). TIAR-1 GFP signal is in red and DNA was counterstained with DAPI (blue). (N=3). (**B**) Single confocal planes of *wild-type* and *tiar-1(tn1543)* fixed two-cell embryos exposed to 34°C for 10 mins (HS) before fixation and immunostained with PQN-59 antibodies (red). DNA was counterstained with DAPI (blue) *(wild-type* n =15 and *tiar-1(tn1543)* n=4, N=1). For both (**A**) and (**B**) enlarged ROIs are shown on the right. Scale bars represent 10 μm.

**S7 Fig.**
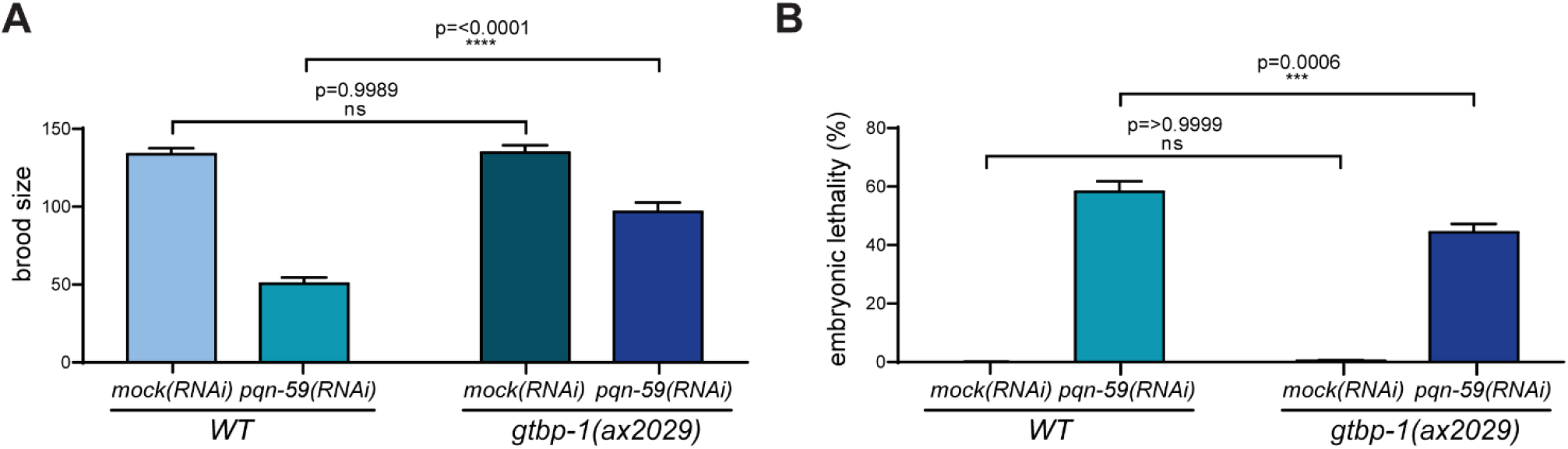
PQN-59 and GTBP-1 co-depletion does not result in an increase of the phenotype of PQN-59 depletion alone. (**A**) Brood size of *wild-type* and *gtbp-1(ax2029)* strains treated with the indicated RNAi. Values correspond to the average number of eggs laid per single worm. The brood size of more than 15 animals of each genotype was evaluated (N=2). (**B**) Embryonic lethality of *wild-type* and *gtbp-1(ax2029)* strains treated with the indicated RNAi. Values correspond to the percentage of un-hatched embryos over the total progeny number (un-hatched embryos and larvae). Embryonic lethality was assessed by counting more than 300 progeny for each condition. N=2. Error bars indicate S.E.M. The P-values were determined using 2way ANOVA test.

**S1 Movie. PQN-59 and GTBP-1 form reversible cytoplasmic granules.** Time lapse movie of *pqn-59::GFP;gtbp-1::RFP* embryos using the CherryTemp temperature-controlled stage. Embryos are recorded at 30°C from minute 00:00 to minute 05:00. From minute 05:00 to minute 20:00 the temperature is 20°C. The temperature shift is visible through the focus change at minute 05:00. Cytoplasmic granules of PQN-59 (green) and GTBP-1 (red) are visible after 5 minutes of heat exposure at 30°C and dissolve after 15 minutes of recovery at 20°C. Scale bar represents 10 μm.

